# Chemical dysbiosis byproducts trigger predation via alternative activation of a peptide quorum sensor in salivarius streptococci

**DOI:** 10.1101/2025.10.09.681396

**Authors:** Guillaume Cerckel, Denis Dereinne, Laura Ledesma-García, Vincent Meuric, Benoît Desguin, Johann Mignolet, Patrice Soumillion, Pascal Hols

## Abstract

Cell-to-cell communication in Gram-positive bacteria is predominantly orchestrated by cytoplasmic sensors of the RRNPPA family. To date, all characterized members of this family are activated by small, unmodified peptide pheromones that mediate bacterial signaling. In the human commensal *Streptococcus salivarius*, the RRNPPA sensor ComR controls both competence (DNA transformation) and predation (bacteriocin production). Here, we reveal that ComR can be dually activated by its cognate peptide (XIP) and a distinct class of small organic molecules. A targeted screen of ∼200 organic compounds identified hydroxyphenylacetic acid (HPAA), a bacterial dysbiosis byproduct accumulating in human fluids, as a potent inducer of ComR. Using *in vivo* and *in vitro* approaches, we demonstrated that HPAA and structurally related carboxylic acids derived from bulky hydrophobic amino acids bind the pheromone-accommodating pocket, leading to ComR activation. Strikingly, while XIP-mediated activation is transient and regulates both competence and predation, HPAA induces a sustained, predation-oriented response. Furthermore, we showed that *Porphyromonas gingivalis*, an oral pathogen, produces sufficient (H)PAA quantity to trigger bacteriocin production in *S. salivarius*, revealing a previously unrecognized chemical interplay between oral microbiota members. These findings highlight the remarkable versatility of cytoplasmic sensors to integrate diverse environmental cues, shedding new light on bacterial peptide-based communication and microbial homeostasis in the human microbiome.

**IMPORTANCE:** Bacterial communication through quorum sensing (QS) is crucial for coordinating key physiological processes. While QS in Gram-positive bacteria has been predominantly associated with peptide pheromones, our study uncovers an undescribed alternative signaling mechanism. We demonstrate that ComR, a cytoplasmic receptor of the RRNPPA family, can be activated by its canonical peptide signal and by small organic molecules derived from the anaerobic breakdown of hydrophobic amino acids. This alternative activation pathway enhances the ability of *S. salivarius* to respond to microbial dysbiosis, maintaining ecological balance in the digestive tract. Beyond revealing a novel layer of bacterial communication, our findings suggest that many RRNPPA-family receptors previously considered “orphan” may respond to yet-undiscovered chemical signals. This work expands our understanding of bacterial sensing and opens new avenues for modulating microbial interactions in health and disease.

## INTRODUCTION

In the complex and competitive environment of the human oral cavity, *Streptococcus salivarius* is a dominant commensal that is recognized for its beneficial role(s) in the homeostasis of this peculiar ecological niche (1–4). The oral cavity is a highly dynamic habitat, occupied by more than 700 bacterial species, which demands adaptive strategies for survival and colonization (3).

*S. salivarius* has developed sophisticated mechanisms to thrive and compete effectively in this challenging environment (1, 3). For instance, its ability to initiate natural transformation, to form biofilm, and its predation behavior through bacteriocin production are of prime importance (2, 5–8). Most of those key processes are regulated by quorum sensing (QS) to synchronize a response at the population level (7–10). However, competence and predation have adverse effects in the short term on cellular metabolism, implying that they are tightly regulated in time and space by local environmental stimuli and stress conditions (8–12).

Two signaling systems regulate competence in streptococci, ComCDE and ComRS (9, 13). Both regulatory pathways rely on the cytosolic production and extracellular export of a genome-encoded small signaling peptide (9). This pheromone is only active after secretion and maturation. The ComCDE system is found in mitis and anginosus groups of streptococci (9, 13) and activated by the <u>c</u>ompetence <u>s</u>ignaling <u>p</u>eptide (CSP) through a phosphorylation cascade between ComD and ComE. The phosphorylated form of the transcriptional regulator ComE will then induce transcription of *comX* (master regulator of competence; alternative sigma factor σ^X^) and late competence genes and, in turn, natural transformation (9, 14, 15). Conversely, the ComRS system found in mutans, bovis, pyogenes, suis, and salivarius streptococcal groups (9, 13) is a direct induction system (16, 17) (Fig. 1A). The pheromone precursor ComS is generally exported through the PptAB ABC transporter and matured by the Eep protease before its external release as a pheromone, named *sig<u>X</u>/com<u>X</u>*-inducing-<u>p</u>eptide (XIP) (11, 18) (Fig. 1A). When it reaches a critical concentration level, XIP will in turn be internalized by the generic oligopeptide transporter Ami/Opp (19). Then, it will bind to the cytoplasmic sensor ComR to form an active ComR**·**XIP complex (17). This complex will induce *comS* to activate a positive feedback loop for the propagation of the signal and *comX* (10, 19) (Fig. 1A). The production of ComX will initiate the late competence phase by recruiting the RNA polymerase to activate the set of genes encoding the transformasome (9) (Fig. 1A).

**Fig. 1.**
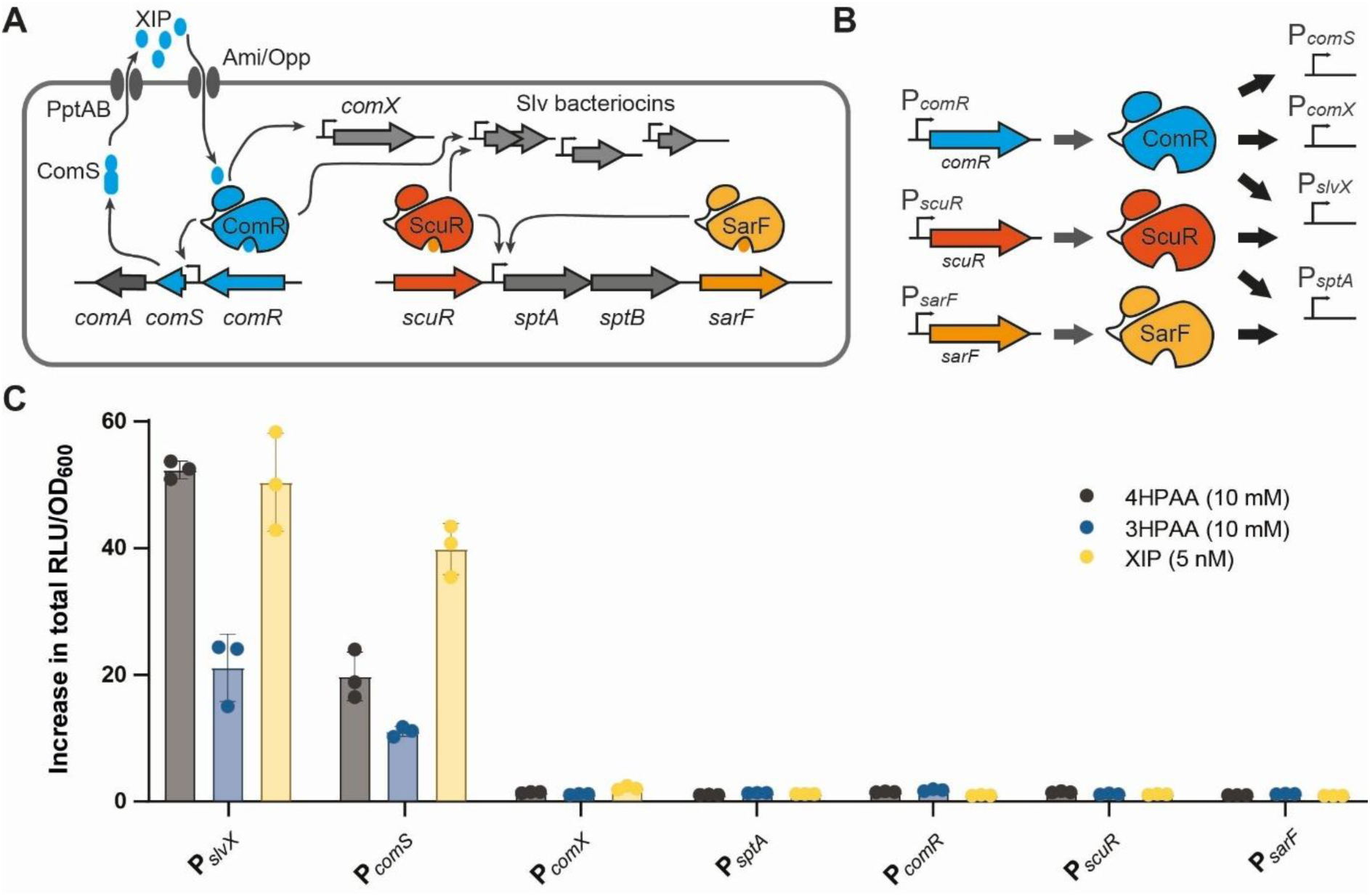
4HPAA and 3HPAA are inducers of predation in *S. salivarius*. (**A**) Scheme illustrating activation of early competence and predation in *S. salivarius*. A basal level of *comS* expression will lead to XIP pheromone accumulation in the medium. When XIP reaches a critical concentration, its reimportation will lead to ComR activation. In turn, it will induce *comS* to amplify XIP production (positive feedback loop) and *comX*, resulting in DNA transformation. It will also coactivate a suite of bacteriocin promoters (*e.g.*, P*_slvX_*) linked to predation. In parallel, activated ScuR is also able to induce bacteriocin genes. In addition, *sptAB,* encoding a bacitracin-like ABC transporter, is induced by both ScuR and SarF. (**B**) For the screen, we used the promoters of *comR*, *scuR*, and *sarF* genes. We also used P*_comX_* (only bound by ComR), P*_slvX_* (bound by ComR and ScuR), and P*_sptA_* (bound by ScuR and SarF). P*_comS_* was not used for the screen but only for validation of *comA* expression (bacteriocin exporter gene). (**C**) Luminescence assays of P*_slvX_*, P*_comS_*, P*_comX_*, P*_sptA_*, P*_comR_*, P*_scuR_*, and P*_sarF_* reporter fusions (*luxAB*) in response to 4-hydroxyphenylacetic acid (4HPAA; 10 mM), 3-hydroxyphenylacetic acid (3HPAA, 10 mM), and XIP (5 nM). Reporter strains were grown in CDMG. The fold increase in total luminescence (RLU/OD_600_) was calculated between induced and non-induced conditions. Data are mean values of biological triplicates (dots) ± standard deviation (error bars).

In addition, predation is co-activated with competence by either ComCDE or ComRS (directly or indirectly) through the production of bacteriocins and lytic enzymes in most streptococci (13). In *S. salivarius*, those two processes are directly coregulated by the ComRS system (8). Besides *comS* and *comX*, the ComR**·**XIP complex directly activates the expression of *comA* that codes for a bacteriocin transporter (operon *comS*-*comA*) and a range of *blp/slv* genes encoding a cocktail of class II bacteriocins (salivaricins) with a variable spectrum of antimicrobial activity (8) (Fig. 1A). Interestingly, expression of bacteriocin genes is more responsive to lower XIP pheromone concentrations than the *comX* gene, suggesting that their production could precede competence activation to induce lysis of neighbor cells and supply exogenous DNA for natural transformation (8). While coupling competence and predation can have advantages, streptococci are also able to uncouple the activation of predation (13, 20). In most streptococci, activation of predation alone is dependent on the QS BlpCRH system, whose activation through phosphotransfer is very similar to the ComCDE system (13, 20). In *S. salivarius*, this system is replaced by a duo of ComR-like regulators, named ScuR and SarF (7). We have shown that the three regulators – *i.e.*, ComR, ScuR, and SarF – recognized a similar DNA palindromic motif but with a different selectivity depending on a slight sequence variation of the binding site (*i.e.*, ComR box) and its relative position in the promoter sequence (7). For instance, ScuR can activate the bacteriocin genes without activating the *comX* gene, allowing the uncoupling of predation activation from competence triggering (7) (Fig. 1A).

While ComR is activated by the XIP pheromone, the native peptide(s) activating ScuR and SarF remain(s) unknown (7). However, we were able to select a range of synthetic activating peptides (sBI7 peptide as prototype) from a mutant library able to selectively activate ScuR/SarF without activating ComR (7). Those three ComR-like regulators belong to a subgroup among the Rgg subfamily of RRNPPA (Rap, Rgg, NprR, PlcR, PgrX, AimR) cytoplasmic sensors, whose mode of activation has been deeply investigated at the biochemical level (21, 22). ComR subfamily members are composed of two domains, the DNA-binding domain (HTH, <u>h</u>elix-turn-<u>h</u>elix) connected by a linker to the peptide-binding domain (TPR, tetratrico<u>p</u>eptide repeats) (17). The TPR domain binds the peptide inside a hydrophobic pocket, whose residues involved in specific interactions with the inducing peptide have been deeply investigated at the structural level (16, 17, 23). The peptide binding induces a range of conformational changes that eventually allow HTH release from TPR interaction, TPR dimerization, and DNA binding (17).

The ComRS and ScuR/SarF systems of *S. salivarius* can only be activated under laboratory conditions by adding their signaling peptides (7, 8). Recently, we showed that the ComRS system of *S. salivarius* is under the strict control of the general stress-sensing system CovRS, indicating that perceiving the right combination of external stimuli from the environment is a key aspect for competence/predation activation (10). Knowing the importance of nutrient availability on the physiology of *S. salivarius* (4, 24, 25), we here explored the impact of a range of carbon/nitrogen sources on the modulation of ComRS and ScuR/SarF systems. From around 200 growth conditions, we unexpectedly identify hydroxyphenylacetic acids as direct inducers of the ComR regulator. With mobility shift assays, modeling and *in vivo* tests, we dissected the chemical moiety responsible for activation and extended the inducers to a family of carboxylic acids derived from hydrophobic amino-acids. Those molecules, naturally produced in case of dysbiosis by anaerobic proteolytic bacteria from the digestive tract (26–28), trigger an unconventional sustained predation response in salivarius streptococci without activating DNA transformation. This dual activation of ComR (cognate pheromone for QS and small organic molecules for dysbiosis sensing) highlights the flexibility of cytoplasmic sensors to respond to multiple stimuli for the maintenance of the microbial homeostasis in the digestive tract.

## RESULTS

### Hydroxyphenylacetic acids trigger predation in *S. salivarius*

Rgg-like systems are highly influenced by external signals and cell physiology. These parameters could either control the expression of the Rgg-encoding genes (29) or affect the formation of the active Rgg-peptide complexes, such as through the modulation of inducing peptide availability or the production of interfering peptides (Fig. 1A) (30–32). To explore the regulation and activation of the three paralogous sensors, ComR, ScuR, and SarF, from *S. salivarius* HSISS4, we designed a screening approach based on six luminescent reporter fusions (Fig. 1B). While a first set of fusions aims to monitor the activation of the promoters of each regulator gene (P*_comR_*, P*_scuR_*, and P*_sarF_*, fused to *luxAB* genes), a second set reports the impact of these regulators on the activation of the downstream genes (Fig. 1A and B). The chosen promoters were (i) P*_comX_* as a proxy of competence that is specifically induced by ComR, (ii) P*_slvX_* as a proxy of predation that is induced by both ComR and ScuR, and (iii) P*_sptA_* controlling a bacitracin-like ABC transporter that is activated by ScuR and SarF (7) (Fig. 1B). Altogether, these six fusions will allow us to monitor any direct or indirect effect of a physiological stimulus that would activate one or more of these three regulatory systems in *S. salivarius*.

Since carbon/nitrogen sources were previously shown to impact the activity of Rgg systems in various streptococci (29, 33), we screened 192 molecules (*e.g.*, simple/complex sugars, amino-acids and derivatives, and organic acids) using phenotype microarrays (Biolog plates PM1 and PM2A) with the six luminescent reporter strains (Table S1 and Fig. S1). As preliminary tests showed limited growth of *S. salivarius* across a wide range of carbon sources, chemically-defined medium (CDM) was supplemented with 0.15% glucose. This concentration of glucose allows an initial growth (OD_600_ ∼0.75), which could be increased if the tested carbon source is sequentially consumed (diauxic growth) or co-metabolized. While no major increase in luminescence was noted for P*_comR_*, P*_scuR_*, P_sar*F*_, P*_comX_*, and P*_sptA_* reporter strains compared to the control (C^-^), two carbon sources induced luminescence with the P*_slvX_* reporter strain (Fig. S1). Interestingly, 4-hydroxyphenylacetic acid (4HPAA) and 3-hydroxyphenylacetic acid (3HPAA) triggered a ∼50- and ∼20-fold upregulation of P*_slvX_*, respectively (Fig. 1C). In addition, P*_slvX_* activation is not due to a higher expression of any of the three regulator genes and does not imply a coactivation of competence, as shown by the absence of P*_comX_* induction (Fig. 1C). Moreover, P*_comS_* was also activated by those two compounds, showing that the bacteriocin exporter gene (*comA*) is co-induced with genes encoding bacteriocins (P*_slvX_* as a proxy) (Fig. 1C).

Together, these results show that predation via the activation of genes encoding bacteriocins and their exporter is stimulated by two small organic acids, 4HPAA and 3HPAA, which are not produced by *S. salivarius* but are known as byproducts or intermediates in the catabolism of aromatic amino acids of many anaerobic bacteria ((34) and KEGG database).

### 4HPAA induces predation through ComR

Since the presence of 4HPAA has been reported as a biomarker of microbial dysbiosis in the digestive tract due to the proliferation of proteolytic bacteria (28), this prompts us to further investigate how 4HPAA stimulates the activation of bacteriocin genes. To confirm the activation of P*_slvX_*, the dynamics of its induction by 4HPAA during growth was compared to the XIP pheromone (Fig. 2A). While XIP addition triggered a sharp activation peak during the early growth phase, as previously reported (8), 4HPAA induced a continuous activation during the exponential growth phase, which is indicative of a different pathway in the perception of the stimulus. We also evaluated if those two modes of activation could be additive, synergistic, or competitive. The kinetics of bacteriocin gene activation by a mixture of a low amount of XIP (5 nM) with 4HPAA (1 mM) display an extended response during growth, resulting from the superposition of XIP and 4HPAA responses (Fig. S2A). We observed that 4HPAA could compete with XIP (∼2-fold decrease of the XIP response) but without major impact on the global level of bacteriocin gene expression (Fig. S2B). Given that P*_slvX_* could be activated by either ComR or ScuR, we examined the impact of 4HPAA on various deletion mutants (Fig. 2B). The deletion of *comR* (Δ*comR*) led to a complete absence of P*_slvX_* induction by 4HPAA. In contrast, the deletion of *comS* (Δ*comS*) did not alter the 4HPAA response, indicating that neither the XIP precursor nor the mature XIP is required for this regulatory pathway. In addition, the deletion of *scuR* and *sarF* genes (Δ*scuR-sarF*) had no impact on 4HPAA stimulation. Altogether, these results show that 4HPAA specifically activates the cytoplasmic sensor ComR.

**Fig. 2.**
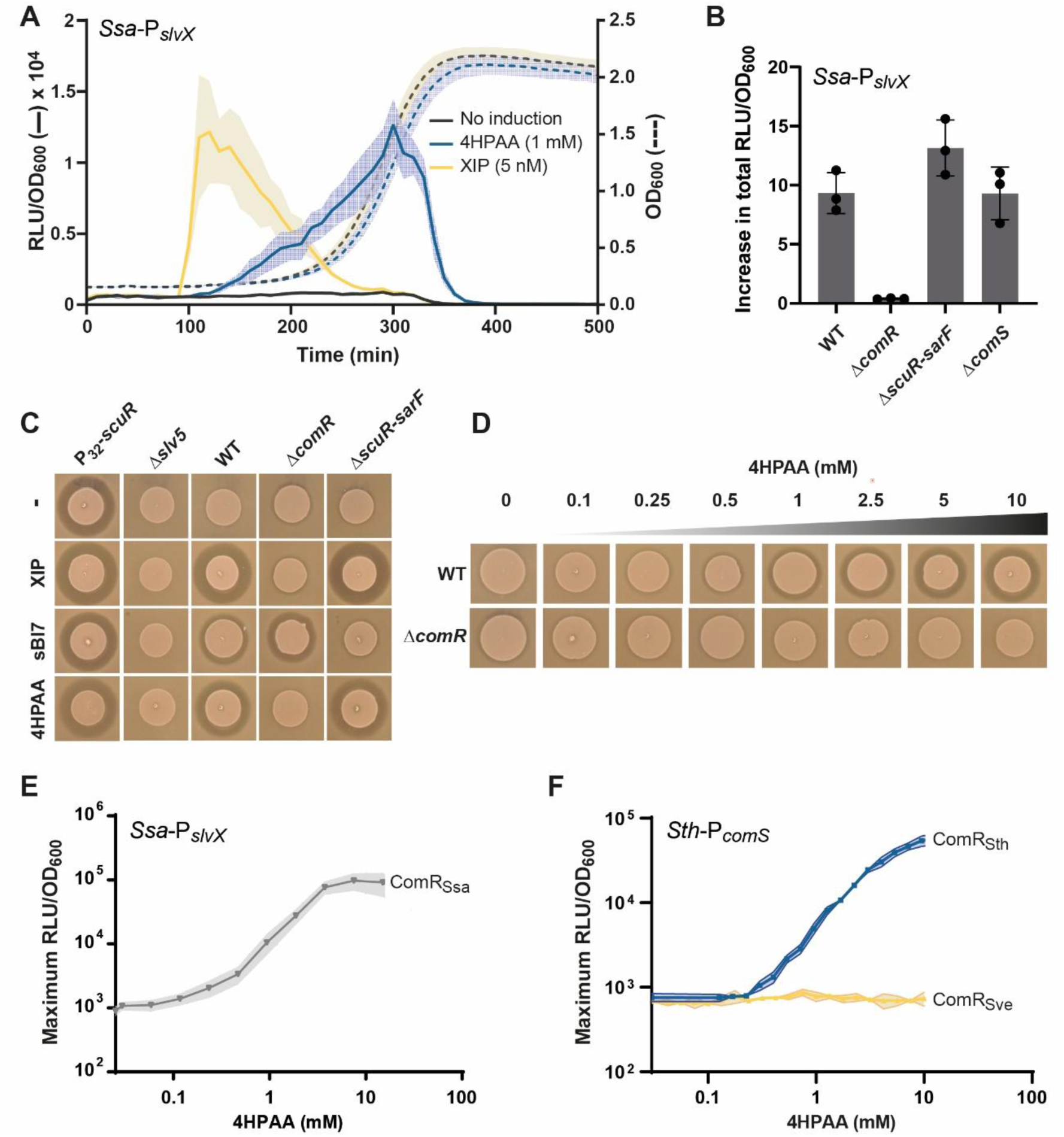
4HPAA activates predation through ComR in salivarius streptococci. (**A**) Luminescence (RLU/OD_600_) over time of the *S. salivarius* P*_slvX_-luxAB* reporter fusion without inducer, with 4HPAA (1 mM), or with XIP (5 nM). (**B**) Fold increase in total luminescence between 4HPAA induction (10 mM) and non-induced conditions for the *S. salivarius* P*_slvX_* reporter strain (WT) and its derivative mutants (Δ*comR*, Δ*scuR-sarF*, and Δ*comS*). (**C**) Bacteriocin assays with wild type (WT) and mutants P*_32_-scuR* (positive control), Δ*slv5* (negative control), Δ*comR*, and Δ*scuR-sarF* without inducer (-), with 1 μM XIP (ComR-specific), 1 μM sBI7 (ScuR/SarF-specific), or 10 mM 4HPAA. *Lactococcus lactis* IL1403 was used as an indicator strain. (**D**) Bacteriocin assays with increasing concentrations of 4HPAA (0 to 10 mM) on wild type (WT) and Δ*comR* mutant (negative control). Indicator strain as in panel C. (**E** and **F**) Maximum luminescence in response to a 4HPAA gradient (0 to 10 mM) for *S. salivarius* P*_slvX_*-*luxAB* (ComR_Ssa_) (**E**), *S. thermophilus* P*_comS_*-*luxAB* Δ*comS* (ComRSth), and its isogenic strain *comR::comR_Sve_* (ComR_Sve_) (**F**). Data presented in panels A, B, E, and F are mean values of biological triplicates (dots in panel B) ± standard deviation (error bars in panel B, and light color zones in panels A, E, and F).

Bacteriocin assays were also performed to confirm that 4HPAA effectively induces a genuine predation response. We tested the wild-type strain, a mutant strain with all five bacteriocin loci deleted (Δ*slv5*; negative control), a mutant constitutively expressing bacteriocins (P_32_*-scuR*; positive control), the Δ*comR* mutant, and the double Δ*scuR*-*sarF* mutant. Four experimental conditions were evaluated: no inducer, addition of XIP (1 µM), sBI7 (1 µM), or 4HPAA (10 mM) (Fig. 2C). With XIP addition, a ComR-specific inducing peptide, both the wild type and the Δ*scuR*-*sarF* mutant produced inhibition halos, but not the Δ*comR* mutant. Conversely, with the addition of the sBI7 peptide, which is ScuR/SarF-specific, the wild type and the Δc*omR* mutant exhibited inhibition halos, but not the Δ*scuR*-*sarF* mutant. Notably, the addition of 4HPAA phenocopies the XIP effect, confirming that the 4HPAA response is strictly ComR-dependent. We also evaluated bacteriocin production in response to an increasing concentration of 4HPAA (Fig. 2D). In the wild type, an inhibition halo was detected at a concentration of 1 mM, and its size increased up to 5 mM. Given that ComR activation by XIP leads to a coupling between predation and competence (8), we also evaluated the capacity of 4HPAA to induce natural DNA transformation in *S. salivarius*. 4HPAA-induced cells (10 mM) failed to generate any detectable transformant in standard conditions (Fig. S2C), corroborating the absence of P*_comX_* activation by 4HPAA (Fig. 1C).

Altogether, these results show that 4HPAA is exclusively perceived by ComR through a pathway that is independent of the XIP pheromone. Moreover, the selective activation of predation via ComR without competence induction is an exclusive feature of the 4HPAA-inducing response.

### 4HPAA induction is ComR-specific among salivarius streptococci

*S. salivarius* (Ssa), *Streptococcus thermophilius* (Sth), and *Streptococcus vestibularis* (Sve) are closely related species forming the salivarius group (1). This is evident from the high similarity among their respective ComR proteins. Indeed, ComR_Ssa_ and ComR_Sth_ display a very high level of identity (95%) and are both activated by the same XIP peptide (XIP_Ssa/Sth,_ LPYFAGCL) (19). ComR_Sve_ is more distantly related (83% of identity with ComR_Ssa_) and is strictly induced by a variant XIP peptide (XIP_Sve_, VPFFMIYY) (16). To test the HPAA response, we used three reporter strains, each encoding a different homolog of ComR: *S. salivarius* P*_slvX_-luxAB* for ComR_Ssa_ (positive control), *S. thermophilus* P*_comS_-luxAB* (Δ*comS*) for ComR_Sth_, and a variant version of the previous strain where ComR_Sth_ was exchanged by ComR_Sve_ (*comR_Sth_::comR_Sve_*) (16). The two isogenic strains of *S. thermophilus* were previously validated for the strict response (P*_comS_* activation) of their ComR version to XIP_Sth_ and XIP_Sve_, respectively (16). We tested a gradient of 4HPAA (0 to 10 mM) on these three reporter strains (Fig. 2E and F). For ComR_Ssa_ and ComR_Sth_ reporter strains, an increase in luminescence started in the range of 100-500 µM with a maximum achieved at ∼8-10 mM. Interestingly, the third reporter strain encoding ComR_Sve_ did not respond to 4HPAA at any tested concentration (Fig. 2F).

Since the more distant ortholog of ComR (ComR_Sve_) is insensitive, these results show that the 4HPAA response is not only ComR-dependent but also ComR-specific, which points towards a direct interaction between 4HPAA (or a derivative compound) and ComR for its activation.

### 4HPAA activates ComR *in vitro*

There was no prior evidence for direct interactions between RRNPPA regulators and organic acids as inducers. Therefore, we were prompted to investigate the direct binding of 4HPAA to ComR *in vitro*. Since ComR_Sth_ and ComR_Ssa_ behave similarly *in vivo*, we used ComR_Sth_ based on its extensive biochemical characterization (16, 17, 23). We performed Electrophoretic Mobility Shift Assays (EMSA) with purified ComR_Sth_ (and ComR_Sve_ used as a negative control), a fluorescent ^Cy3^P*_comS_* DNA probe, and 4HPAA (or XIP_Sth_ used as a positive control) as inducer of complex assembly (16, 17) (Fig. 3A and Fig. S3). In these experiments, we added 4HPAA at a single concentration directly into the running buffer to ensure a homogenous distribution of the molecule. We used gradients of ComR_Sth_ and ComR_Sve_ (noting that an excess of ComR leads to its precipitation) along with a fixed amount of the DNA probe and performed migration in buffers with and without 4HPAA (10 mM). As a positive control, we also used conditions with a fixed concentration of ComR_Sth_ and a gradient of XIP_Sth_ to visualize the position of the ComR**·**XIP complex. Notably, ComR_Sth_ in the presence of 4HPAA led to the formation of a specific complex with similar mobility compared to the XIP_Sth_ positive control (Fig. 3A). Conversely, ComR_Sve_, which does not respond to 4HPAA *in vivo* (Fig. 2D), displayed no high molecular size complex (Fig. S3).

**Fig. 3:**
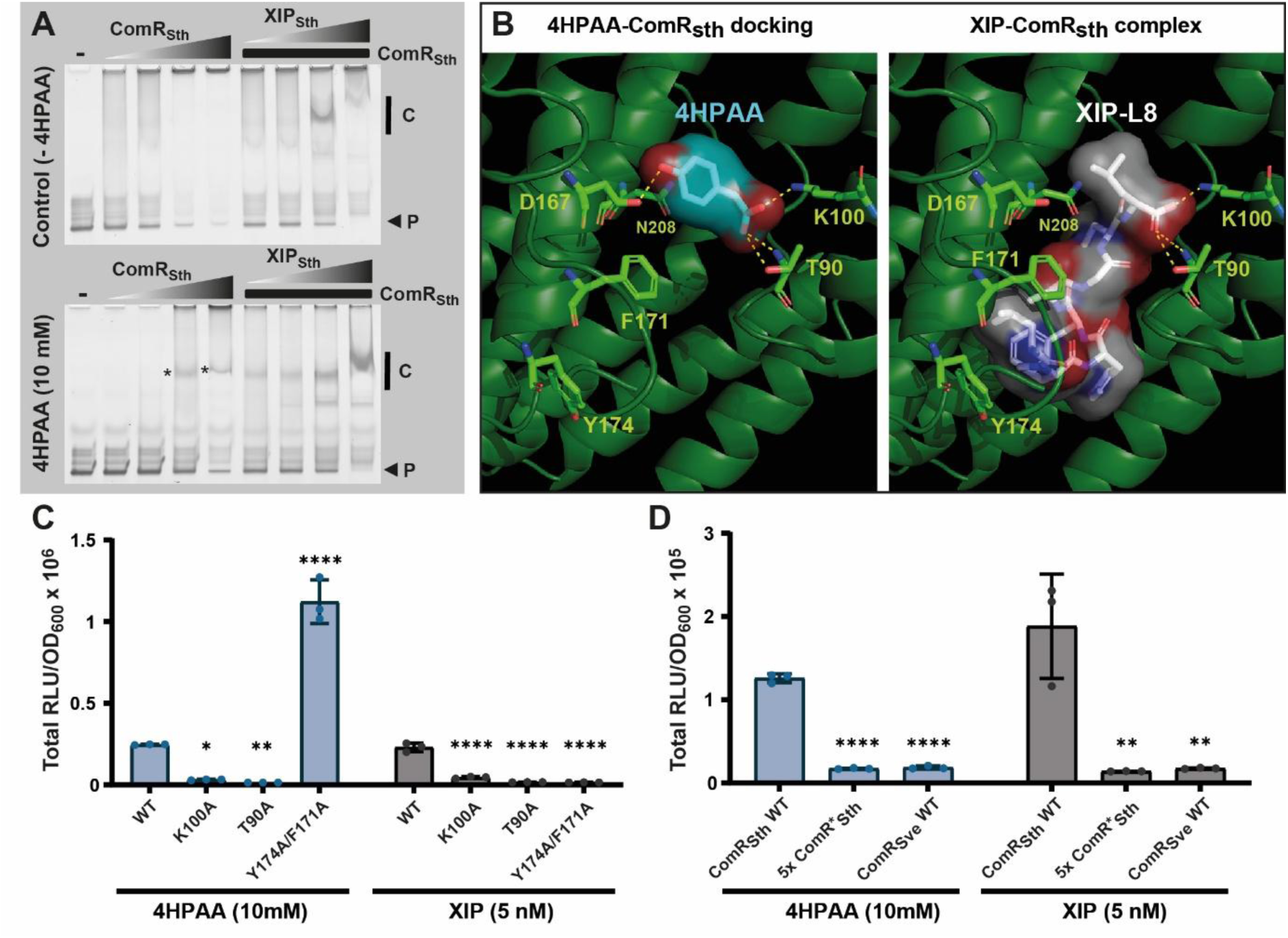
Direct binding of 4HPAA into the peptide pocket of ComR. (**A**) Mobility shift assays of the *comS* promoter probe (40 ng) conducted with a gradient of purified ComR_Sth_ (gray triangles, 2:2 dilutions from 4 μM) (left) or with a fixed concentration of ComR_Sth_ (2 μM) and a gradient of XIP_Sth_ (gray triangles, 25, 50, 200, and 1000 nM) (right), without (top) or with (bottom) 10 mM 4HPAA. Probes (P) indicated by a black arrow are Cy3-conjugated DNA fragments of 40 bp. Vertical back lines indicate the position of specific complexes and stars label complexes formed in the presence of 4HPAA alone. The control condition without ComR_Sth_ is indicated with a minus sign (-). (**B**) *In silico* prediction of 4HPAA in complex with ComR_Sth_ holoform (right) compared to the native XIP-ComR_Sth_ complex (PDB 5JUB) (left). The top-ranked docked 4HPAA (blue) is mimicking the positioning of XIP-L8 (white). ComR residues T90, K100, and D167 are highlighted in light green, as they were predicted to directly interact with 4HPAA (yellow dotted lines). ComR residues F171 and Y174 are shown since they are essential for the interaction with XIP. (**C**) *In vivo* luminescence response (total RLU/OD_600_) of ComR_Sth_ wild-type (WT) and variants K100A, T90A, and Y174A/F171A to 4HPAA (10 mM) or XIP (5 nM). Experiments were performed with *S. thermophilus* P*_comS_*-*luxAB* Δ*comS*. (**D**) *In vivo* luminescence response of ComR_Sth_ wild-type (WT), ComR_Sth_ penta-mutant (5x ComR*_Sth_) responding only to XIP_Sve_, and comR_Sve_ WT to 4HPAA (10 mM) or XIP (5 nM). Same genetic background as in panel C. Dots, bars, and error bars in panels C and D show biological triplicates, mean values, and standard deviations, respectively. All variants were statistically compared to ComR_Sth_ WT using one-way ANOVA with Dunnett’s test (*, *P* < 0.05; **, *P* < 0.01; ****, *P* < 0.0001).

Together, these results showed the direct binding of 4HPAA to ComR, resulting in the formation of an active complex able to bind DNA. To the best of our knowledge, a chemical compound has never been reported to directly activate a regulator of the RRNPPA family.

### 4HPAA is predicted to occupy the XIP C-terminus binding site for ComR activation

To investigate how 4HPAA could interact with ComR, we run *in silico* docking predictions using the EADock DSS engine from SwissDock (35). We used the anionic form of 4HPAA and compared docking predictions for the holoforms (XIP-activated) of ComR_Sth_ and ComR_Sve_, and the apoform of ComR_Sth_. Remarkably, the top-ranked prediction (total energy of –17.7 kcal/mol) for holo-ComR_Sth_ positioned the 4HPAA molecule deeply in the XIP binding pocket right at the site occupied by the pheromone C-terminal leucine (XIP-L8) (Fig. 3B). By comparison, the predicted docking of 4HPAA in the same configuration within the XIP-binding pocket of holo-ComR_Sve_ is less stable (total energy of -14.8 kcal/mol), and such positioning is not predicted in apo-ComR_Sth_. The carboxylate group of docked 4HPAA is predicted to form a salt bridge with the side chain ammonium group of residue K100 from helix α7 and to H-bond with T90 from loop α6-α7 of ComR_Sth_, as previously reported for the C-terminal carboxylate group of XIP-L8 (Fig. 3B) (17). In addition, the hydroxyl group of 4HPAA is predicted to H-bond with D167 from helix α10, which could enhance its stabilization in the pocket (Fig. 3B). Finally, N208 is contacting the 4HPAA aromatic ring in an adequate orientation to establish a weak H-bond with the π cloud as an acceptor (NH…π bond) (36).

We previously reported that mutating residues at the bottom (T90, K100; interaction with XIP-L8) or the top (F171/Y174; interaction with XIP-L1) of the XIP binding pocket in ComR_Sth_ abolished the XIP_Sth_-mediated activation (Fig. 3B) (17, 23). Mutants T90A and K100A lost their ability to induce luminescence in response to 4HPAA, mirroring our previous observations with XIP (17) (Fig. 3C). In contrast, the double Y174A/F171A mutant exhibited a 4-fold increase in light emission upon 4HPAA activation compared to the control, despite a loss of inducibility by XIP (Fig. 3C). These effects are partially recapitulated in the single F171A mutant (Fig. S4). The F171A and Y174A/F171A mutants were previously shown to display a higher basal level of activation without XIP addition (23), indicating a less stringent conformational control that could explain their better activation by the remote binding of 4HPAA. In a previous work, we also generated a ComR_Sth_ variant (5x ComR*) with five substitutions (R92G, V205A, S248G, S289K, and I290T) that responds exclusively to XIP_Sve_ due to a remodeling of the peptide binding pocket (16). Our luminescence assays showed that, as observed with ComR_Sve_, this ComR_Sth_ variant is unable to respond to 4HPAA (Fig. 3D). A dissection of individual substitutions in the XIP binding pocket showed that the R92G substitution recapitulates the loss of 4HPAA induction, as previously reported for its key role in the XIP_Sth_-mediated activation mechanism (17) (Fig. S4). In addition, the S289K substitution showed an interesting behavior, being more reactive to 4HPAA and less sensitive to XIP_Sth_ than their respective controls (Fig. S4). The lysine at position 289 could favor initial encountering with 4HPAA to facilitate its activation effect.

Overall, this analysis supports a model where 4HPAA partially mimics the XIP activation mechanism by substituting XIP-L8 for key interactions with residues T90 and K100 in the peptide binding pocket of ComR. This alternative mode of activation is further supported by the 4HPAA ability to activate ComR mutants that are not or are weakly responding to XIP.

### Carboxylic acid derivatives of bulky hydrophobic amino acids are ComR inducers

In our investigation of 4HPAA selectivity for ComR activation, we tested several structural variants of this molecule for their *in vivo* activation capacity with a specific focus on three key aspects: (i) the carboxylate function and its associated negative charge, (ii) the polarity around the aromatic ring, and (iii) the overall size of the molecule (Fig. 4A). For variations of the carboxylate moiety, we examined tyramine (featuring a positively charged ammonium at neutral pH) and methyl-4HPAA (a neutral methylester derivative), but none of them elicited a luminescence signal (Fig. 4B). This aligns with our 4HPAA-binding prediction, which suggests that the carboxylate is critical for interactions with residues T90 and K100 inside the XIP binding pocket (Fig. 3B). In addition, two α-hydroxy-carboxylates, (R)-(-)-mandelic acid and D-(+)-3-phenyllactic acid, also failed to induce luminescence (Fig. 4). This is in line with the structural model that predicts steric hindrance upon substituting the pro-R α-hydrogen by a bulkier group or extending the length of the carboxylate moiety. For the polarity around the aromatic ring, phenylacetic acid (PAA), which lacks a hydroxyl group, or compounds with similar steric hindrance, like 3HPAA and 3-fluoro-4HPAA (Fig. 4), were able to produce luminescence, although at ∼2-fold lower levels (orange bars, Fig. 4B). This indicates that the presence, position, or polarity of the hydroxyl group plays a secondary role in ComR activation, suggesting that 4HPAA interaction with residue D167 is not critical. Furthermore, pentafluoro-PAA failed to induce luminescence (Fig. 4B), further supporting the potential NH…π H-bond stabilization since fluorine atoms will decrease the electron density of the π cloud (36). We also examined compounds with a larger steric hindrance than 4HPAA. The presence of one or two methoxy groups in the meta position of the aromatic ring, as in homovanillic acid and 4-hydroxy-3,5-dimethoxy-PAA, or the addition of a second aromatic ring, as in 1-naphthylacetic acid, resulted in inactive compounds (Fig. 4).

**Fig. 4.**
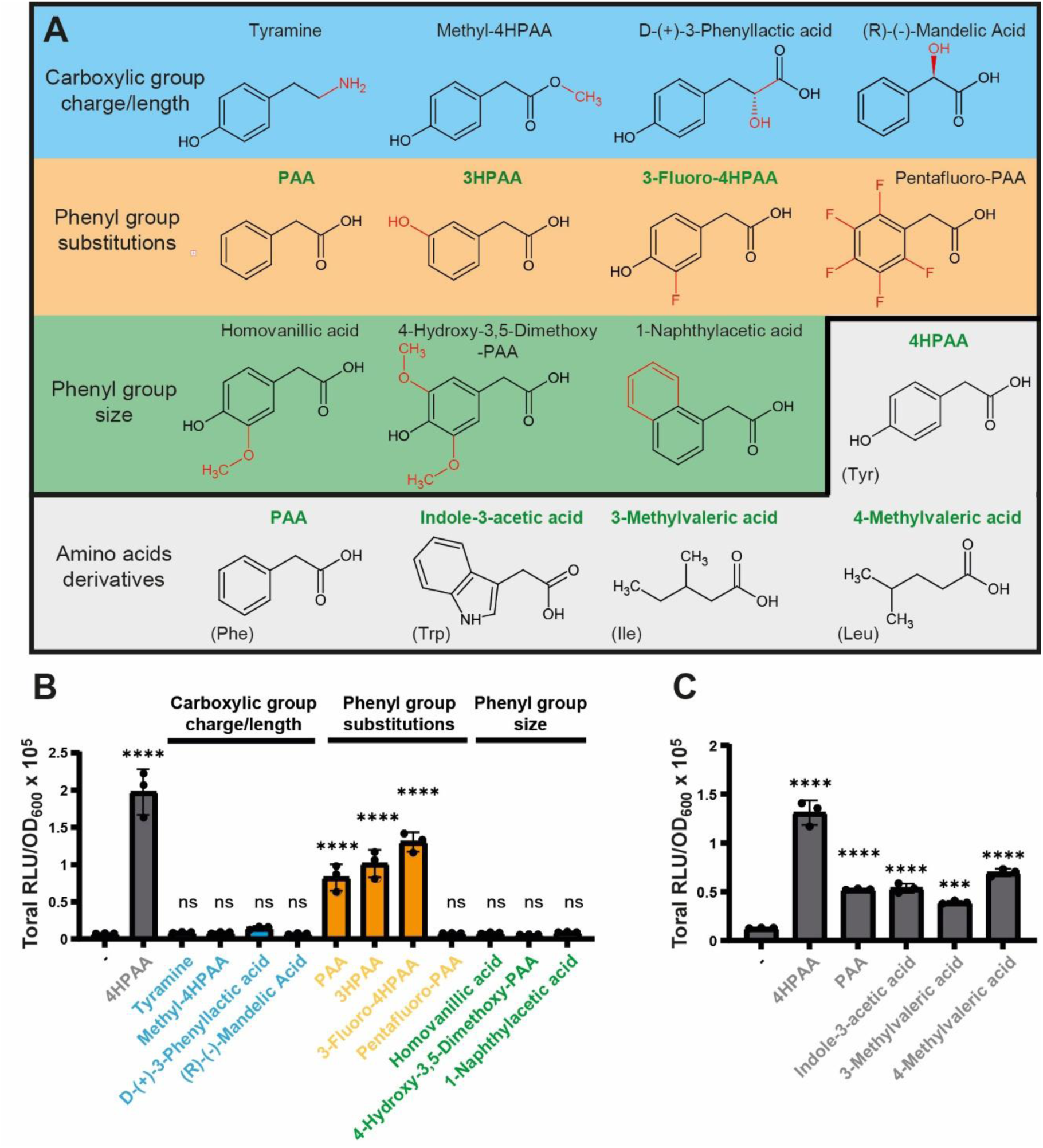
Extended activation of ComR by bulky derivatives of hydrophobic amino acids. (**A**) Structure of 4HPAA variants with modifications of the carboxylic moiety (charge and length; blue rectangle), phenyl group substitutions (orange rectangle), steric hindrance (green rectangle); and hydrophobic amino acid derivatives (gray). Modified chemical groups compared to 4HPAA are highlighted in red. The names of chemicals in bold green indicate inducing molecules. (**B**) Luminescence assays (total RLU/OD_600_) with 4HPAA structural variants (10 mM) shown in panel A. (**C**) Luminescence assays with organic acids derived from bulky hydrophobic amino acids (10 mM) shown in panel A. Derivatives of Tyr, Phe, Trp, Ile, and Leu induced a signal. In panels B and C, experiments were performed with *S. thermophilus* P*_comS_*-*luxAB* Δ*comS*. Dots, bars, and error bars show biological triplicates, mean values, and standard deviations, respectively. All variant molecules were statistically compared to the control condition without organic acid (minus sign) using one-way ANOVA with Dunnett’s test (***, *P* < 0.001; ****, *P* < 0.0001; ns, non-significant).

Since 4HPAA and PAA are two carboxylic acids derived from aromatic amino acids (Tyr and Phe), we also tested equivalent derivatives from the whole set of hydrophobic amino acids for ComR_Ssa/Sth_ activation (Fig. 4C and Fig. S5). In addition to 4HPAA and PAA, the bulkiest compounds, indole-3-acetic acid (IAA; Trp derivative), 3-methylvaleric acid (3MVA, Ile derivative), and 4-methylvaleric acid (4MVA, Leu derivative), were also able to increase the luminescence signal (Fig. 4C), while isovaleric acid (Val derivative) with one carbon less than MVA and other amino acid derivatives of shorter length did not (Fig. S5). Among these three activating compounds, we choose 4MVA for EMSAs, which confirmed its ability to activate ComR as observed for 4HPAA (Fig. S3).

Together, these *in vivo* assays with 4HPAA structural variants strengthen the docking model of ComR activation, regarding (i) the key role played by the carboxylate group and hydrophobic side chain, (ii) the simulating effect of the hydroxyl group in the para position of the aromatic ring, and (iii) the size fitting of the molecule into the binding pocket. Moreover, we pinpoint five amino acid derivatives that could be of biological relevance as signaling molecules since most of them were previously identified in various human fluids associated with microbial dysbiosis (26–28, 37–40).

### *Porphyromonas gingivalis* culture supernatant triggers ComR activation

*S. salivarius* is a dominant commensal species of the oral cavity and the upper part of the small intestine (1, 25). In cases of microbial dysbiosis, the proliferation of Gram-negative bacteria (*e.g.*, *Prevotella* sp. and *Porphyromonas* sp.) in the oral cavity or Gram-positive bacteria (*e.g.*, *Clostridium* sp.) in the small intestine was reported to incompletely catabolize aromatic amino acids, leading to increased levels of 4HPAA, PAA, or IAA in saliva, urine, or feces (28). To evaluate if one of these bacteria could accumulate a substantial level of those compounds in their culture supernatants, we tested the well-studied opportunistic pathogen *P. gingivalis* of the oral cavity (41). *P. gingivalis* W83 was grown under anaerobic conditions in enriched BHI medium (BHIe) until late stationary growth. Filtered supernatants were then added (25% v/v of CDM) to cultures of *S. thermophilus* and *S. salivarius* reporter strains (P*_comS_-luxAB* and P*_slvX_*-*luxAB*, respectively) for luminescence assays (Fig. 5A). The *S. thermophilus* reporter strain was strongly activated by *P. gingivalis* culture supernatants (unextracted *Pg* SN, Fig. 5A), while the BHIe medium alone was inefficient to stimulate light emission. In addition, culture supernatants of *P. gingivalis* grown in BHIe enriched with Phe or Tyr significantly increased the luminescence signal (∼3-fold), supporting the hypothesis that byproducts of aromatic amino-acid catabolism contribute to activation (Fig. S6A). We also validated that the supernatant effect was ComR-dependent and ComS/XIP-independent by using Δ*comR* and Δ*comS* mutants of the reporter strain, respectively (Fig. 5B). To exclude the presence of an inducing peptide in the culture supernatant that could mimic XIP, we also used an oligopeptide transporter mutant (Δ*ami/opp*), which remains similarly inducible as the control strain (Fig. 5B). Intriguingly, the *S. salivarius* reporter strain did not respond in the same conditions, except for a weak activation with aromatic amino-acid enriched BHIe medium (Fig. S6B). We identified that the BHIe medium alone has a quenching effect on the activation of the ComRS system through a downregulation of *comR* expression (P*_comR_*-*luxAB*) (Fig. S6C). To counteract this effect, we used an isogenic reporter strain containing a P*_xyl2_*-*comR* cassette (xylose-inducible) (10). At a low *comR* expression level (xylose 0.2%), the *S. salivarius* reporter strain was activated by the different culture supernatants as shown above with *S. thermophilus* (Fig. S6D).

**Fig. 5.**
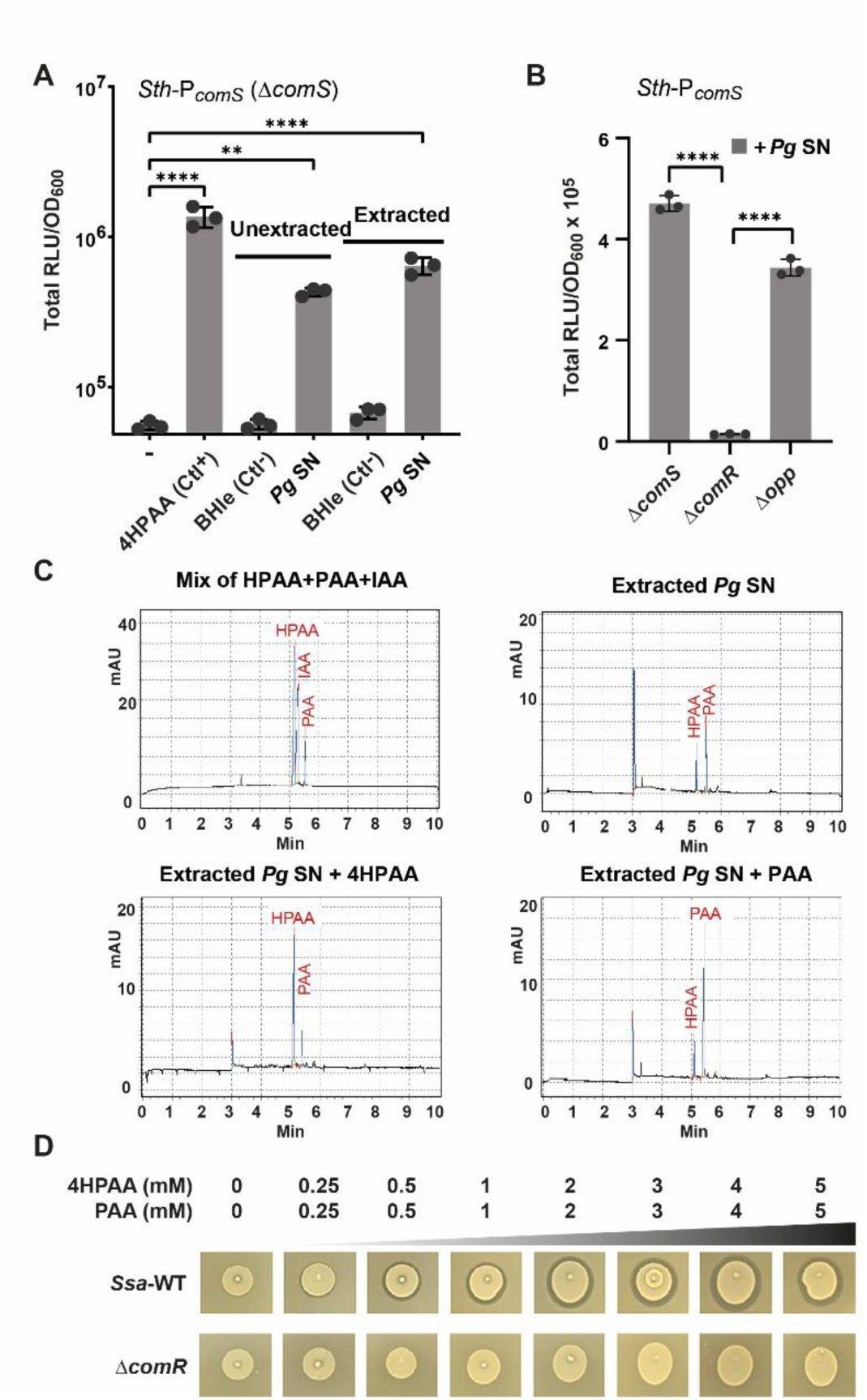
*P. gingivalis* culture supernatant triggers ComR activation. (**A**) Luminescence assays (total RLU/OD_600_) without addition (-), with 1 mM 4HPAA (positive control), and 25% (vol/vol) of BHIe medium (negative control), filtered *P. gingivalis* supernatant (*Pg* SN), ethyl acetate-extracted BHIe (negative control), and ethyl acetate-extracted *Pg* SN in CDM. Experiments were performed with *S. thermophilus* P*_comS_*-*luxAB* Δ*comS* (**B**) Luminescence assays (total RLU/OD_600_) of Δ*comS*, Δ*comR* and Δ*opp* mutants in presence of 25% (vol/vol) of *Pg* SN in CDM. The three mutants are derivatives of *S. thermophilus* P*_comS_*-*luxAB*. (**C**) Separation by capillary electrophoresis of ethyl acetate-extracted *P. gingivalis* supernatant. Electro-chromatograms display a mixture of 4HPAA (3 mM), IAA (2 mM), and PAA (1 mM) used as standards (top left); extracted *Pg* SN (top right); addition of 0.5 mM 4HPAA to extracted *Pg* SN (bottom left); and addition of 0.5 mM PAA to extracted *Pg* SN (bottom right). (**D**) Bacteriocin assays with increasing concentrations of 4HPAA and PAA (each from 0 to 5 mM) on *S. salivarius* wild type (WT) and Δ*comR* mutant (negative control). *Lactococcus lactis* IL1403 was used as an indicator strain. In panels A and B, dots, bars, and error bars show biological triplicates, mean values, and standard deviations, respectively. In panel A, all tested conditions were statistically compared to the control condition without external addition (minus sign). In panel B, the Δ*comS* and Δ*opp* mutants were statistically compared to the Δ*comR* mutant. Both statistical analyses used a one-way ANOVA with Dunnett’s test (**, *P* < 0.01; ****, *P* < 0.0001).

Since these results converged towards the presence of aromatic amino-acid derivatives in the supernatant, we performed an ethyl acetate extraction at low pH to selectively extract hydrophobic acids such as HPAA, PAA, and IAA (42) for their analysis by capillary electrophoresis. We showed that the extracted fraction was still able to generate a specific luminescence signal (extracted *Pg* SN, Fig. 5A) and appeared to contain a mixture (∼1:1) of HPAA and PAA, as shown by electro-chromatograms with appropriate standards (Fig. 5C). Assuming that those two compounds are the main inducing molecules in the unextracted supernatant of *P. gingivalis*, we evaluated their individual concentration to be in the range of ∼4-5 mM based on a dose-response curve with equimolar concentrations of HPAA and PAA (Fig. S6E). To validate that a 1:1 mixture of HPAA and PAA can induce bacteriocin production in *S. salivarius* wild type, bacteriocin assays were performed with increasing concentrations of both compounds. Remarkably, growth inhibition was detected at a ∼10-fold lower concentration than found in the *P. gingivalis* filtered supernatant (Fig. 5D).

Together, these results show that the well-known amino acid degrader, *P. gingivalis* (43, 44), secretes enough aromatic amino acid derivatives in the extracellular growth medium to activate predation in salivarius streptococci.

## DISCUSSION

Cell-to-cell communication in *Bacillota* (low-GC Gram-positive bacteria) is largely controlled by the RRNPPA family of cytoplasmic QS receptors (∼ 5,000 members identified so far) (21). Those sensors are involved in a range of key physiological processes such as horizontal gene transfer, metabolism, virulence, sporulation, predation, biofilm formation, and phage lysogeny (21, 22). So far, all the characterized regulators of this family were reported to be activated by small unmodified peptides used as communication pheromones (22). Notably, we report here that a member of the RRNPPA family could alternate activation by binding its cognate signaling peptide or a specific class of small organic molecules.

In streptococci, RRNPPA members mostly belong to the Rgg family that includes the ComR group of competence-predation signaling systems (21, 45). *S. salivarius* is equipped with three members of this group that coordinate the coupling (via ComR) or the uncoupling (via ScuR/SarF) of predation with competence for DNA transformation (7, 8). In this work, we show an unexpected uncoupling mechanism of predation via ComR by the sensing of byproducts from hydrophobic amino-acid catabolism (named hereafter HACs for <u>h</u>ydrophobic <u>a</u>mino-acids <u>c</u>atabolites) produced by many proteolytic bacteria proliferating in the digestive tract (28) (Fig. 6). This parallel layer of predation control by small organic molecules has substantial advantages compared to the classical regulatory cascade of the ComR-XIP system. In its QS mode for intra-species communication, ComRS is quickly shut down by the intracellular degradation of the XIP pheromone (ComX-PepF relay) to avoid an over-activation of competence that could be deleterious for the cell (11). This mode of control results in a pulse production of bacteriocins (Fig. 2A and 6A), which are expected to release extracellular DNA from neighboring bacteria for DNA transformation (fratricide and sobrinicide processes) (6). In its HAC-responding mode, the ComRS system is not shut down and continuously expresses bacteriocin during growth until the late stages of the exponential phase (Fig. 2A and 6B). This continuous production at high cell density would have the huge benefit of accumulating high concentrations of antimicrobial peptides in the ecological niche of *S. salivarius*. Another advantage of responding to small organic molecules is their higher stability compared to signaling peptides, which could be prone to degradation/sequestration in the extracellular medium by a range of QS quenching mechanisms. Interestingly, HACs largely accumulate in human fluids in cases of microbial dysbiosis in the digestive tract, suggesting quite high stability (26–28, 39, 40). To conclude, those two modes of ComRS activation not only increase the range of activating stimuli but also improve the capacity of predation in time (no shut-down) and space (unlinked to pheromone availability), an asset for a dominant species in various parts of the digestive tract.

**Fig. 6.**
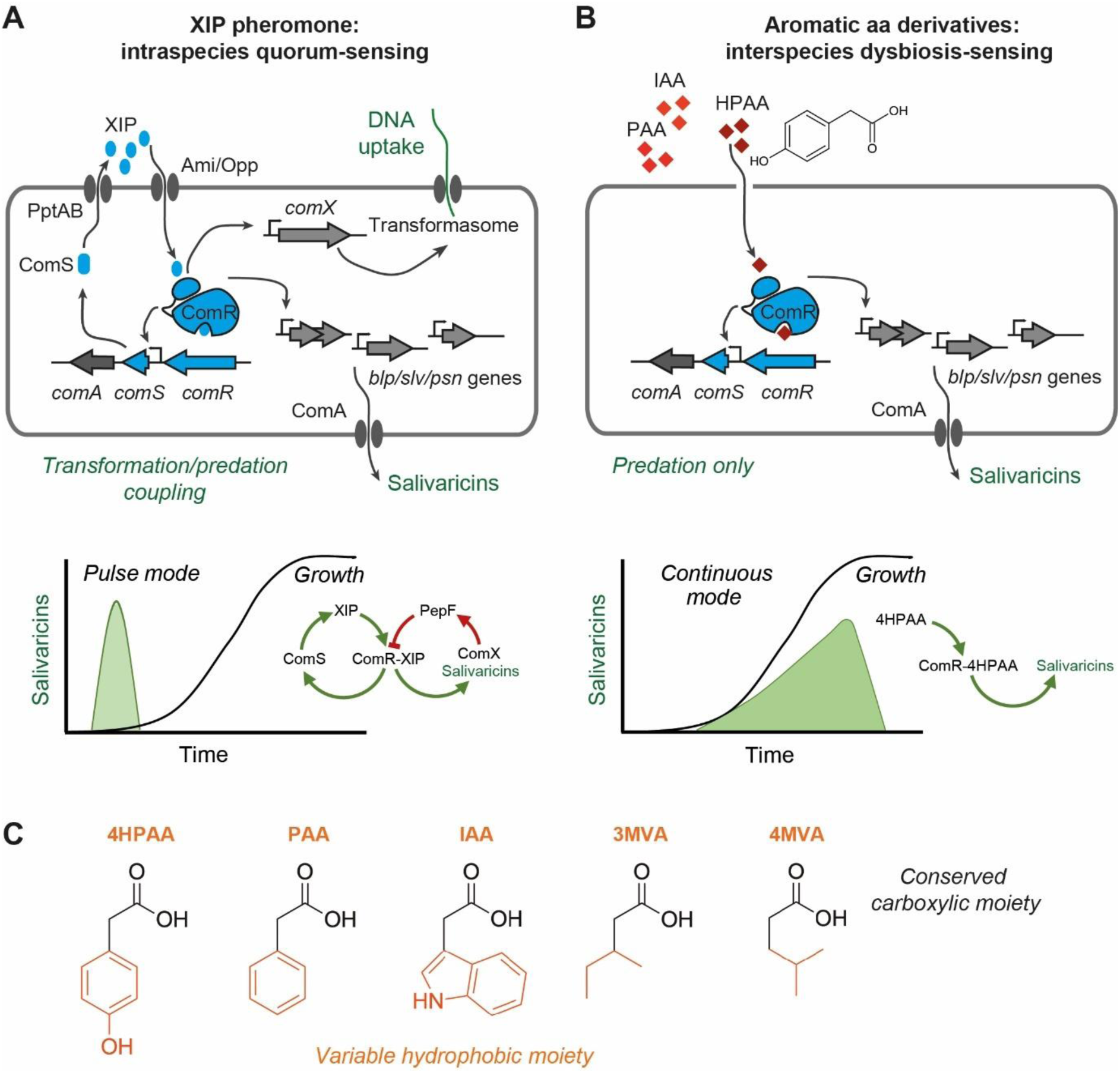
Dual activation of the cytoplasmic sensor ComR by XIP pheromone or aromatic amino acid catabolites in salivarius streptococci. (**A**) Classical activation of the ComRS system by XIP for intraspecies QS, leading to a coactivation of competence (DNA uptake) and predation (salvaricin production). Production of salivaricins takes place in a pulse mode due to a combined positive and negative feedback loop through ComS and the ComX-PepF relay (XIP degradation), respectively. (**B**) Pheromone-independent activation of ComR by aromatic amino acid catabolites for interspecies dysbiosis-sensing, leading only to the activation of predation. Production of salivaricins takes place in a continuous mode due to an absence of a negative feedback loop. (**C**) Summary of ComR-inducing carboxylic acids comprising a conserved carboxylate moiety and a hydrophobic group, which are derivatives of bulky hydrophobic amino-acids.

Previously, we showed that ComR_Sth_ activation by XIP required key interactions at both the bottom (XIP-L8 with ComR-T92 and K100) and the top (XIP-L1 with ComR-Y174 and F171) of the binding pocket (Fig. 3B) (17, 23). Therefore, the ComR apo-form is double-locked and needs interactions at both sub-sites to be activated by XIP (23). Based on *in silico* docking, the use of ComR mutants, and a range of HAC analogs, we propose that HACs activate ComR through a lower stabilization of the active holo-form by binding to the bottom sub-site only. The deep part of the XIP-binding pocket in the apo-form is occupied by a network of six interacting water molecules (two are in direct interaction with T90 and D167) (Fig. S7A). As classically proposed for hydrophobic ligands, HAC binding will release those trapped water molecules (46) while being stabilized through ionic and H-bond interactions with key ComR residues (mainly K100, T90, and D167) of the bottom sub-site (Fig. 3B). We showed that the size of the binding pocket combined with these specific interactions restricts the range of ligands to a couple of activating molecules (*i.e.*, 4HPAA, 3HPAA, PAA, IAA, 3MVA, and 4MVA), all sharing a carboxylic group associated with a hydrophobic moiety of the appropriate bulkiness (Fig. 6C). This model of activation by HACs is strongly supported by the ComR_Sth_ Y174A-F171A variant with a mutated top sub-site, which is insensitive to XIP but strongly activated by the best inducer, 4HPAA. The absence of activation of ComR from *S. vestibularis* (and the mutant ComR_Sth_-R92G) could be explained by a larger binding pocket (Fig. S4 and Fig. S7B) that does not fit as well to 4HPAA (Fig. S7B), whose interaction may not be sufficient to displace trapped water molecules. Moreover, the intermediate level of ComR_Sth_ activation by HACs compared to XIP is probably the reason why HACs are unable to activate competence through transcriptional activation of P*_comX_*. Indeed, we previously reported that the transcription of *comX* was less responsive to XIP activation than bacteriocin genes due to a non-canonical ComR-binding box (8). Hence, although the ComR**·**HAC complex is less stabilized than ComR**·**XIP, this level of stabilization may be sufficient to support higher-order complex formation at the P*_slvX_* promoter but not at P*_comX_*. Altogether, these two levels of control on a unique sensor allowed an uncoupling between predation activation and competence development.

Aromatic HACs have previously been reported as signaling molecules in different bacterial processes such as plant-bacteria symbiosis (47), virulence shutdown (48), and phase variation control for virulence activation (49). Among these processes, the molecular mechanism of signaling regulation by HCAs has only been elucidated for invasion through phase variation in *Neisseria meningitidis*, a colonizer of the nasopharynx (49) that is also occupied by *S. salivarius* (50). In this case, the process is under the control of the stand-alone transcriptional repressor NadR (MarR family), which is directly activated by 4HPAA binding (49, 51). Here, we propose that HACs could act as signaling molecules of dysbiosis resulting from the proliferation of proteolytic bacteria capable of partial degradation of aromatic amino acids. The production of aromatic HACs by bacteria proliferating in the digestive tract has previously been reported for a range of anaerobic Gram-negative and -positive bacteria such as the pathobionts *P. gingivalis* (43) and *Clostridioides difficile* (34, 52), respectively. In addition, aco-occurrence between their proliferation in the case of dysbiosis and the detection of aromatic HACs in human fluids has been largely documented (26–28, 39, 40). As an example, up to 0.1 mM of 4HPAA has been detected in the saliva of patients suffering from periodontal disease (40), a concentration that could be much higher at the site of production. In laboratory conditions, we showed that the periodontal pathogen *P. gingivalis* can produce a mixture of 4HPAA and PAA (mM range) that largely exceeds the minimal concentration required to activate predation in *S. salivarius* (Fig. 6). Although the cocktail of bacteriocins produced by the *S. salivarius* strain used in this study did not affect *P. gingivalis* growth in laboratory conditions, it could inhibit *Streptococcus gordonii*, a key mutualist partner of *P. gingivalis* in multi-species communities (Fig. S6F) (53). Interestingly, a recent study showed that lytic bacteriophages targeting *S. gordonii* in model biofilms with *P. gingivalis* were indirectly capable of inhibiting *P. gingivalis* proliferation (54). So, activation of *S. salivarius* predation by HACs could reshape bacterial communities towards homeostasis through direct or indirect effects on pathobionts.

In conclusion, this study describes an unexpected mode of activation of *S. salivarius* ComR, known as a signaling-peptide cytoplasmic sensor, by a specific class of small organic molecules derived from the anaerobic catabolism of bulky hydrophobic amino acids. This signaling mode may increase the sensory capacity of this beneficial commensal to initiate predation in response to dysbiosis in the human digestive tract. Because many RRNPPA members are pheromone orphans, this research opens a new avenue in chemical communication for the discovery of novel inducing molecules for *Bacillota* cytoplasmic sensors. Our results also pave the way for innovative therapeutic strategies where small hydrophobic acids could stimulate bacteriocin production in commensal bacteria to regulate/inhibit pathogen populations.

## MATERIAL & METHODS

### Carbon/nitrogen sources, organic acids, and synthetic peptides

Carbon/nitrogen sources from Biolog plates PM1 and PM2A are listed in Table S1. Organic acids and their suppliers are listed in Table S2. All organic acids were diluted in water to 100 mM and neutralized with an equimolar amount of NaOH. Synthetic octapeptides XIP_Sth_ (LPYFAGCL) and sBI7 (LPFWLILG) (purity of 95%) and salivaricins BlpK, SlvV, SlvW, SlvX, SlvY, SlvZ, and PsnL (6) were supplied by Peptide 2.0 Inc. (Chantilly, VA, USA) and resuspended in 100% dimethyl sulfoxide (DMSO). Final concentration was quantified using a Nanodrop apparatus (ThermoFisher Scientific).

### Bacterial strains, plasmids, and oligonucleotides

Bacterial strains, plasmids, and oligonucleotides used in this study are listed and described in supplementary information (Tables S3 and S4).

### Growth conditions

*Escherichia coli* TOP10 (Invitrogen) were cultivated with shaking at 37°C in LB (Lysogeny Broth). *S. salivarius* HSISS4 and *S. thermophilus* LMD-9 derivatives were grown at 37°C without shaking in M17 (Difco Laboratories, Detroit, MI) or in chemically-defined medium (CDM) (55) supplemented with 1% (w/v) glucose (M17G, CDMG, respectively), except when specified. The Sodium β-glycerophosphate (19 g/l) buffer used in M17 was also added to CDM to increase growth and luminescence induction. *Lactococcus lactis* IL1403 was grown in M17G at 30°C without shaking. *S. gordonii* LMG17843 was grown in liquid M17G or M17G agar plates supplemented with 5% (v/v) defibrinated horse blood (bioMérieux, France) at 37°C. Agar 1.5% (w/v) was added into M17 and LB plates. Solid plates inoculated with *S. salivarius* cells were incubated anaerobically in jars (Oxoid AnaeroGen 2.5 L, Thermo Fischer Scientific) at 37°C. Ampicillin (250 µg.ml^−1^), spectinomycin (200 µg.ml^−1^), chloramphenicol (5 µg.ml^−1^), or erythromycin (10 µg.ml^−1^) were added as required. *P. gingivalis* W83 (ATCC308) cultures were restarted from frozen samples on Columbia agar plates supplemented with yeast extract (5 g.l^-1^), defibrinated horse blood (5% v/v), hemin (25 mg.l^−1^), and menadione (10 mg.l^−1^) at 37°C under a 5% CO_2_ atmosphere for 2-3 days. Afterwards, colonies were transferred into liquid Brain Heart Infusion medium supplemented with yeast extract (5 g.l^-1^), hemin (25 mg.l^−1^), and menadione (10 mg.l^−1^) (BHI_e_) and incubated for 3-5 days. When needed, 15 mM tyrosine or 15 mM phenylalanine was added to BHIe to stimulate the catabolism of aromatic amino acids. Culture supernatants of cells grown until an OD_600_ of ∼ 1.9 were collected by centrifugation (5,000 × *g* for 15 min) and then filtered (0.22 µm). This supernatant was kept at -80°C before its addition to reporter strain cultures.

### Construction of mutants and reporter strains

Null mutants were constructed by exchanging (double homologous recombination) the coding sequences (CDS) of target genes (sequence between start and stop codons) for an erythromycin resistance cassette using natural transformation as reported before (8). Integration of the antibiotic resistance cassette at the right location was subsequently checked by PCR. The promoters of *rgg* genes were fused to the *luxAB* reporter genes and inserted with a chloramphenicol resistance cassette at the permissive tRNA threonine locus (*HSISS4_r00061*) by double homologous recombination. All DNA fragments (Table S5) were amplified by PCR using the Phusion high-fidelity polymerase (Thermo Fischer Scientific) following a protocol as recommended by the manufacturer. Overlapping PCR products were transferred in competence-induced HSISS4 derivatives, *erm* and *luxAB*-*cat* cassettes were amplified from pGIUD0855*ery* and pJIM*cat*, respectively.

### Luciferase assays

Overnight precultures were diluted at a final OD_600_ of 0.05. Culture samples (300 μl) were incubated in the wells of a sterile covered black microplate with a transparent bottom (Greiner Bio-One, Alphen a/d Rijn, the Netherlands). Growth (OD_600_) and luciferase (Lux) activity (expressed in relative light units, RLU) were monitored at 10-min intervals during 24 h in a Hidex Sense multimode reader (Hidex, Turku, Finland). Total or maximum specific Lux activity was obtained by dividing Lux activity by the OD_600_ for each measurement and summing all the data obtained over time or selecting the maximum value, respectively. Experimental values represent the averages of at least three independent biological replicates. Statistical analyses of multiple comparisons to the control mean were performed with one-way ANOVA with Dunnett’s test. Standard deviations and *P* values were calculated.

For the screening of carbon/nitrogen sources using Phenotype Microarrays (Biolog plates PM1 and PM2A), commercial plates were unsuitable for luminescence assays. The carbon/nitrogen sources from the original plates were initially dissolved in 100 µl of CDM without glucose and incubated for 10 min at 37°C with agitation to ensure complete dissolution. Subsequently, this solution was transferred to the wells of a sterile black microplate with a transparent bottom. A preculture of the reporter strain was added to obtain a final OD_600_ of ∼ 0.05. Glucose from a 25% (w/v) stock solution was added to achieve a final concentration of 0.15% (v/v), and the total volume in each well was adjusted to 300 µl with additional CDM. Growth and luminescence were then monitored as reported above.

### Bacteriocin assays

The spot-on-lawn (multilayer) detection method (7) used to test bacteriocin production by *S. salivarius* was performed as follows. Overnight cultures of producer strains were diluted in fresh M17G medium and grown until mid-log phase (OD_600_ of ∼0.5). In parallel, we casted plates with a bottom feeding layer (M17G, 1.5% agar) supplemented with inducing molecules (XIP_Sth_, sBI7, 4HPAA, or 4HPAA + PAA in equimolar ratio), when required. Next, an overnight culture of *L. lactis* IL1403, used as an indicator strain, was incorporated in pre-warmed soft M17G medium (0.3% agar) at a final OD_600_ of 0.05 and casted as a top layer. Finally, we spotted 3 μl of the producer strains on the top layer. Plates were incubated overnight in anaerobic conditions before analysis of the inhibition zones surrounding the producer colonies.

A variation of the above procedure was applied to test the activity of chemically-synthesized salivaricins against *P. gingivalis* W83 and *S. gordonii* LMG17843. Stationary-phase cultures (2-3 days) of *P. gingivalis* W83 or diluted overnight cultures (OD_600_ of 0.05) of *S. gordonii* LMG17843 were spread evenly with sterile cotton swabs on their respective solid media using the lawn method. Onto these lawns, 2 μl of synthetic bacteriocin (100 μM) was spotted. Plates were incubated overnight under appropriate growth conditions until lawns were established, after which inhibition zones at the application sites were assessed.

### Natural DNA transformation assays

*S. salivarius* overnight precultures in CDMG were diluted in the same medium at a final OD_600_ of 0.05. After an incubation of 75 min at 37°C, 4HPAA (10 mM) or XIP_Sth_ (5, 200, or 500 nM) and PCR-amplified DNA fragments containing an *erm* gene (Δ*scuR*::*erm*) surrounded by 1.5-kb recombination arms (1 μg) were added. Cells were grown for 4 h at 37°C before plating on selective M17G agar and incubation in anaerobic conditions.

### ComR purification

*E. coli* TOP10 strains containing plasmids pBAD*comR_Sth_*-*streptag* (56) and pBAD*comR_Sve_*-*streptag* (16) were used for the purification of ComR_Sth_-StreptagII and ComR_Sve_-StreptagII proteins, respectively. Protein purification and storage were performed as previously described (16). Protein purity was analyzed by SDS-PAGE and protein concentration was measured using a Nanodrop apparatus (Thermo Fisher Scientific).

### Electrophoretic mobility shift assays

EMSA assays were performed as previously described (7, 16). Twofold serial dilutions of purified ComR protein (initial concentration of 4 μM) were mixed with a 40-bp dsDNA fragment (40 ng) carrying the ComR box of P*_comS_* coupled to the Cy3 fluorophore in the binding buffer. In parallel, a fixed concentration of purified ComR protein (2 μM) was mixed with a gradient dilution of XIP_Sth_ (5, 50, 200, and 1000 nM) together with ^Cy3^P*_comS_* (40 ng). Negative controls were performed in the absence of protein. Samples were incubated at 37°C for 10 min prior to separation on a native 4 to 20% gradient gel (iD PAGE gel; Eurogentec). Carboxylic acids were added to the running buffer (MOPS) at a final concentration of 10 mM. DNA complexes were detected by fluorescence on the Amersham Typhoon biomolecular imager (Cytiva) with bandpass excitation and emission filters of 595/25 nm (Cy3). The double-stranded DNA fragment was obtained from annealing of single-stranded Cy3-labeled (at the 5’ end) and unlabeled oligonucleotides.

### Supernatant extraction and analysis by capillary electrophoresis

Considering the water solubility of 4HPAA at different pHs and its solubility in ethyl acetate (42), a dedicated extraction protocol was developed to extract 4HPAA and closely related molecules from *P. gingivalis* culture supernatants. Initially, 4 ml of culture supernatant was acidified using 0.1 ml of concentrated hydrochloric acid (HCl, 37% w/w). Subsequently, 2 ml of ethyl acetate was incorporated into the solution, followed by thorough mixing and a 10-min decantation period. The organic phase was then separated and mixed with a sodium hydroxide (NaOH) solution (2 ml, 0.01 M). After a further 10-min period of decantation, the aqueous phase was isolated, neutralized with HCl (37% w/w), and filtered (0.22 µm).

For analytical characterization, the extracted sample was separated using a capillary electrophoresis apparatus (Capel 105M, Lumex Instruments), with boric acid (0.1 M, pH ∼ 8.5) as the background electrolyte. A bare fused silica capillary of 54/46 cm in total/effective length with an internal diameter of 50 μm from Agilent was used. Samples were injected at a pressure of 30 mbar for 5 s. A voltage of 25 kV was applied throughout the analysis. UV absorbance at 210 nm was used for detection. The capillary temperature was maintained at 20°C during all steps. The analyses were conducted on the sample alone and in the presence of either 0.5 mM 4HPAA or 0.5 mM PAA, allowing for the assessment of the extraction efficiency and the selectivity of the protocol towards 4HPAA and closely related molecules.

### 4HPAA docking and structure prediction or visualization

For *in silico* docking predictions of 4HPAA in ComR, we used the EADock DSS engine from SwissDock (35). Docking was performed with the crystal structure of 4HPAA (CCDC 274674) in its anionic form. The holoforms of ComR_Sth_ (PDB 5JUB, chain A) and ComR_Sve_ (PDB 6HUA, chain A), and the apoform of ComR_Sth_ (PDB 5JUF, without SO_4_ and H_2_O molecules) were used as targets. Structure predictions of ComR mutants with AlphaFold 2 were obtained from the AlphaFold CoLab notebook (57). The figures with structural elements were prepared by using the graphic software PyMol (http://www.pymol.org/).

## AUTHOR’S CONTRIBUTION

GC, LLG, PS, and PH conceived and designed the study. GC, DD, VM, and BD carried out the laboratory work. GC, DD, LLG, BD, PS, and PH analyzed and interpreted the data. GC, VM, BD, JM, PS, and PH wrote or revised the manuscript. All authors read and approved the final manuscript.

## COMPETING INTERESTS

LLG, JM, and PH declare that they are listed as inventors on patent(s) or patent application(s) related to bacteriocin production and uses.

## ACKNOWLEDGMENTS

The work of PH was supported by the Belgian National Fund for Scientific Research (FNRS, grants PDR T.0110.18/T.0111.22 and CDR J.0090.21) and the Concerted Research Actions (ARC, grants 17/22-084 and 22/27-120) from Federation Wallonia-Brussels. GC held a doctoral fellowship from FNRS (FRIA fellowship). JM received funding from the European Union’s Horizon 2020 research and innovation program (Marie Skłodowska-Curie grant N° 101018461). PH and BD are, respectively, Research Director and Research Associate at the FNRS.

## SUPPLEMENTAL INFORMATION

Supplemental information includes 5 tables, 7 figures, and 2 datasets.

## DATA AVAILABILITY

All data are contained within the manuscript. All numerical data and statistical analyses are provided as supplemental dataset 1. All original images used in figures are provided as supplemental dataset 2.

