## Supplementary material for "Chemical dysbiosis byproducts trigger predation via alternative activation of a peptide quorum sensor in salivarius streptococci": Fig. S1, Fig. S2, Fig. S3, Fig. S4, Fig. S5, Fig. S6, Fig. S7, Table S1, Table S2, Table S3, Table S4, Table S5

**CONTENT:**

**SUPPLEMENTARY FIGURES**

**Fig. S1. Screening of carbon/nitrogen sources using Phenotype Microarrays.**

**Fig. S2. Impact of XIP and/or 4HPAA on predation and competence in *S. salivarius*.**

**Fig. S3. Mobility shift assays to assess direct binding of different organic acids.**

**Fig. S4. *In vivo* testing of all available ComR$_{Sth}$ mutants involved in XIP$_{Sth}$ binding or selectivity change towards XIP$_{Sve}$.**

**Fig. S5. Luminescence assays with hydrophobic amino acids and their organic acid derivatives.**

**Fig. S6. Luminescence assays with culture supernatants of *P. gingivalis* and growth inhibition by salvaricins.**

**Fig. S7. Mechanism of ComR$_{Sth}$ activation by 4HPAA.**

**SUPPLEMENTAL TABLES**

**Table S1. List of tested carbon/nitrogen sources and light emission of reporter strains from Biolog plates PM1 and PM2A.**

**Table S2. List of organic acids and amino acids tested in this study.**

**Table S3. List of strains and plasmids.**

**Table S4. List of oligonucleotides used in this study.**

**Table S5. Overlapping and cloning PCR sub-fragments**

### SUPPLEMENTAL FIGURES

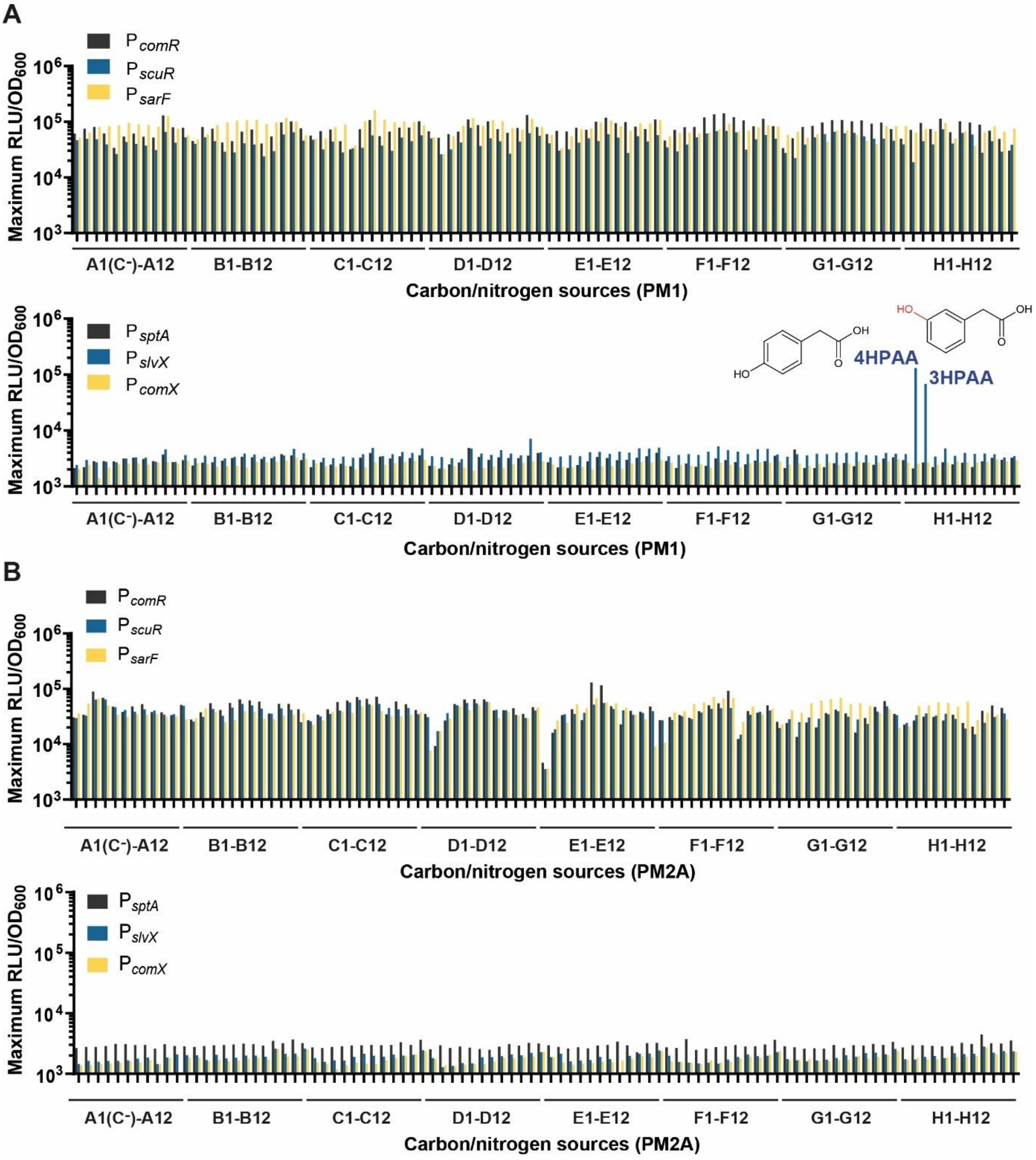

**Fig. S1. Screening of carbon/nitrogen sources using Phenotype Microarrays.** Data were obtained with Biolog plates PM1 (**A**) and PM2A (**B**). $P_{comR}$, $P_{scuR}$, $P_{sarF}$, $P_{sptA}$, $P_{slvX}$, and $P_{comX}$ reporter fusions (*luxAB*) were tested for maximum light emission (RLU/$OD_{600}$) in CDM supplemented with 0.15% glucose. Both 4-hydroxyphenylacetic acid (4HPAA) and 3-hydroxyphenylacetic acid (3HPAA) increased luminescence of $P_{slvX}$ but none of the other promoters. Carbon/nitrogen sources and luminescence data from Biolog plates PM1 and PM2A are listed in Table S1.

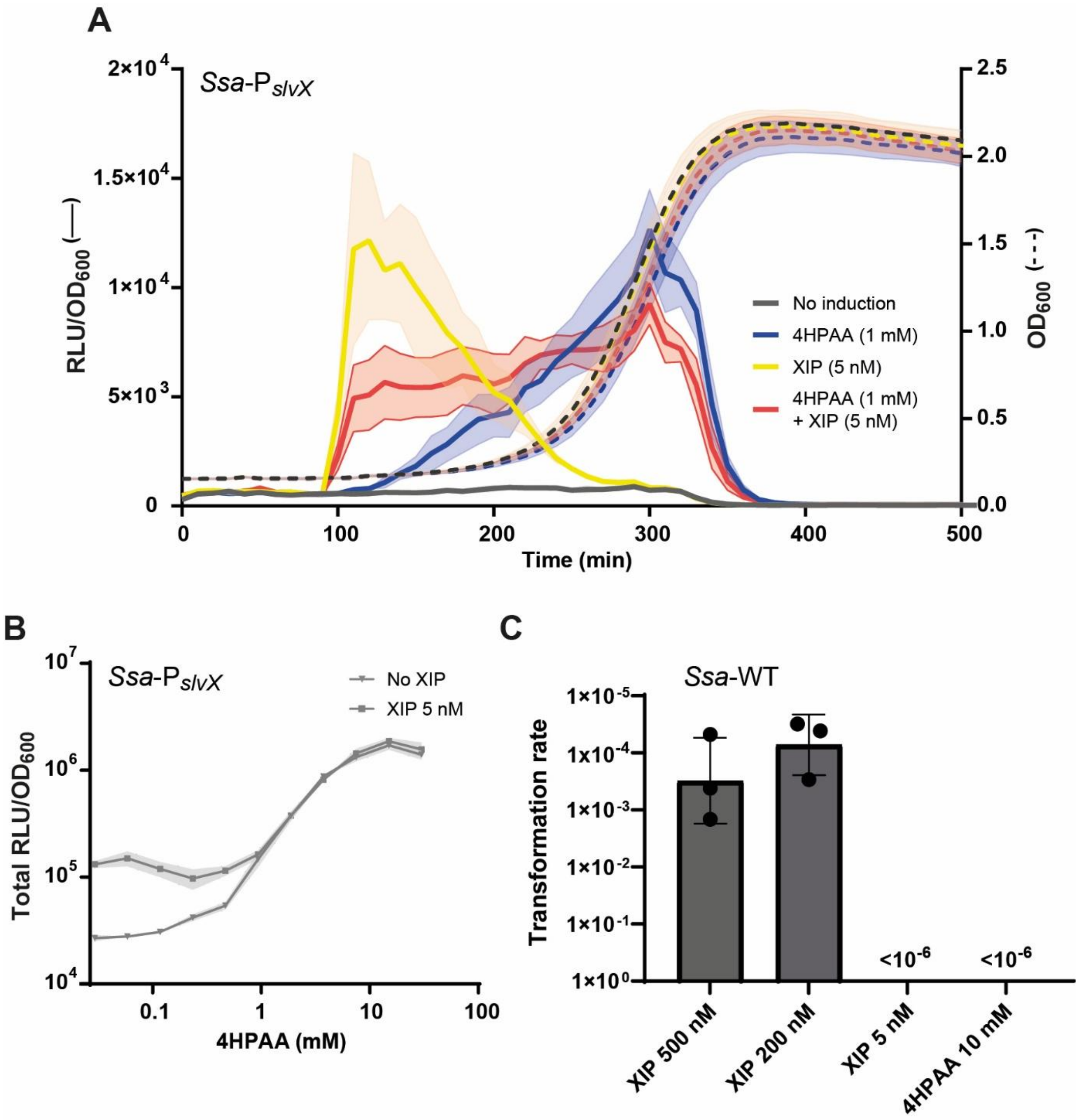

**Fig. S2. Impact of XIP and/or 4HPAA on predation and competence in *S. salivarius*.** (**A**) Luminescence (RLU/$OD_{600}$) over time of the *S. salivarius* $P_{slvX}$-*luxAB* reporter fusion without inducer, with 4HPAA (1 mM), XIP (5 nM), or 4HPAA (1 mM) + XIP (5 nM). (**B**) Total luminescence (RLU/$OD_{600}$) in response to a 4HPAA gradient (0 to 30 mM) in absence (No induction) or addition of XIP (5 nM) for *S. salivarius* $P_{slvX}$-*luxAB*. (**C**) *S. salivarius* HSISS4 (WT) transformation assays with XIP (5, 200, and 500 nM) or 4HPAA (10 mM), reported to 1 µg/ml. Data are mean values of biological triplicates ± standard deviation (light color zones in panels A and B).

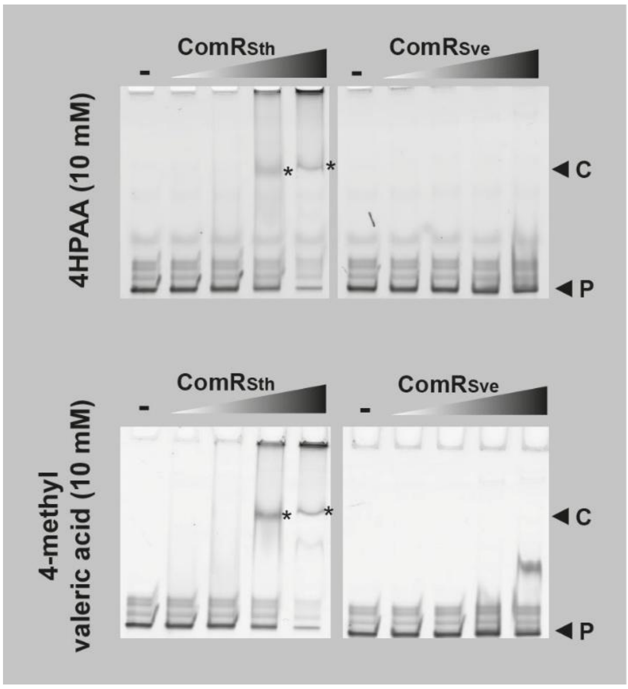

**Fig. S3. Mobility shift assays to assess direct binding of different organic acids.** Mobility shift assays of the *comS* promoter probe (40 ng) conducted with gradients of purified $ComR_{Sth}$ (left) and $ComR_{Sve}$ (right) (Gray triangles, 2:2 dilutions from 4 µM) with 10 mM 4HPAA or 4-methylvaleric acid in the running buffer. Black arrows with a P label are the positions of the probes consisting of Cy3-conjugated DNA fragments of 40 bp. Black arrows with a C label are the position of specific complexes and stars label complexes formed with $ComR_{Sth}$ in presence of organic acids that are not formed with $ComR_{Sve}$ (negative control). The control condition without ComR is indicated with a minus sign (-). For each organic acid, the two parts correspond to samples that were run on the same gel.

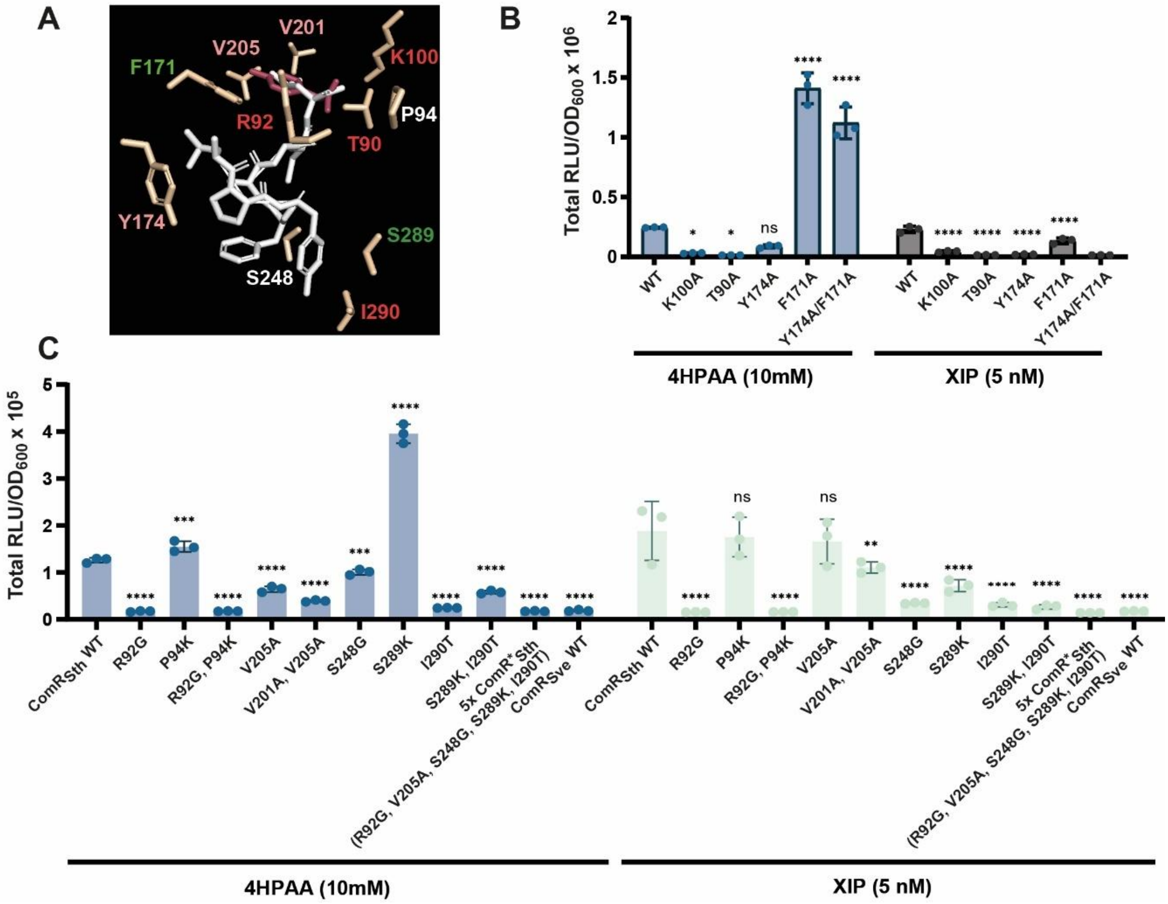

**Fig. S4.** ***In vivo* testing of all available ComR$_{Sth}$ mutants involved in XIP$_{Sth}$ binding or selectivity change towards XIP$_{Sve}$.** (**A**) Positioning of the mutated ComR$_{Sth}$ residues in the XIP$_{Sth}$ binding pocket. Mutated residues strongly or partially affecting 4HPAA induction are labelled in red and pink, respectively. Mutated residues showing enhanced 4HPAA activation are labelled in green. Mutated residues, which are neural regarding 4HPAA induction, are labelled in white. XIP$_{Sth}$ and predicted 4HPAA docking are shown in white and burgundy red, respectively. (**B**) *In vivo* luminescence response (total RLU/OD$_{600}$) of ComR$_{Sth}$ wild-type (WT) and variants K100A, T90A, Y174A, F171A, and Y174A/F171A to 4HPAA (10mM) or XIP (5 nM). Experiments were performed with *S. thermophilus* P$_{comS}$-*luxAB* Δ*comS*. (**C**) *In vivo* luminescence response of ComR$_{Sth}$ wild-type (WT), ComR$_{Sth}$ penta-mutant (5x ComR*$_{Sth}$), comR$_{Sve}$ WT, and variants R92G, P94K, R92A/P94K, V205A, V201A/V205A, S248G, I290T to 4HPAA (10 mM) or XIP (5 nM). The genetic background used as in panel B. In panels B and C, dots, bars, and error bars show biological triplicates, mean values, and standard deviations, respectively. All variants were statistically compared to ComR$_{Sth}$ WT using one-way ANOVA with Dunnett's test (*, $P < 0.05$; ***, $P < 0.001$; ****, $P < 0.0001$; ns, non-significant).

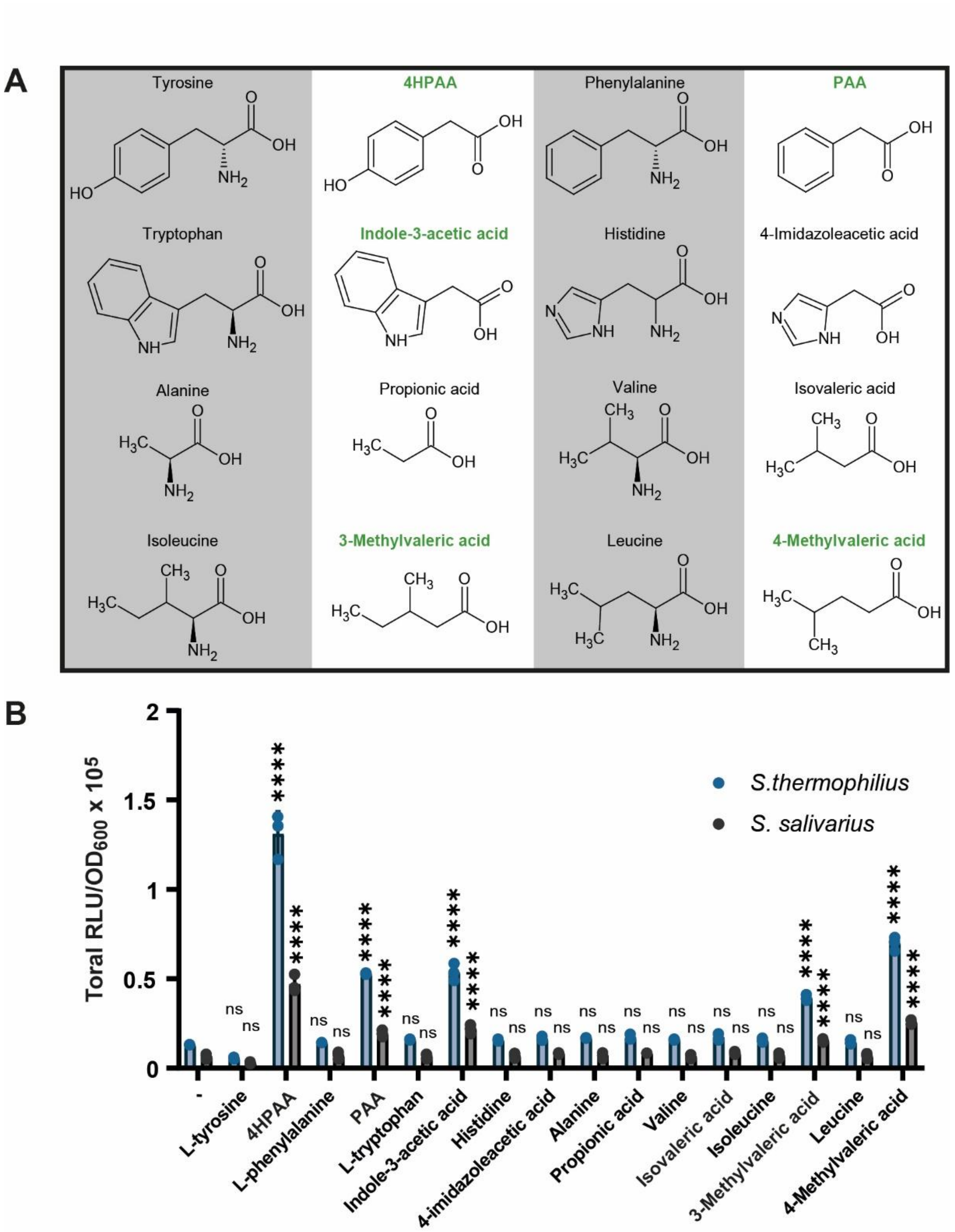

**Fig. S5. Luminescence assays with hydrophobic amino acids and their organic acid derivatives.** (**A**) Structure of hydrophobic amino acids and their organic acid derivatives. The name of chemicals in bold green indicates inducing molecules. (**B**) Luminescence assays (total RLU/$OD_{600}$) with hydrophobic amino acids and their derivatives (10 mM) shown in panel A. Derivatives of Tyr, Phe, Trp, Ile, and Leu induced a signal. Experiments were performed with *S. thermophilus* $P_{comS}$-*luxAB* Δ*comS* and *S. salivarius* $P_{slvX}$-*luxAB*. Dots, bars, and error bars show biological triplicates, mean values, and standard deviations, respectively. All variant molecules were statistically compared to their respective control condition without addition of amino acid or organic acid derivative (minus sign) using one-way ANOVA with Dunnett's test (****, $P < 0.0001$; ns, non-significant).

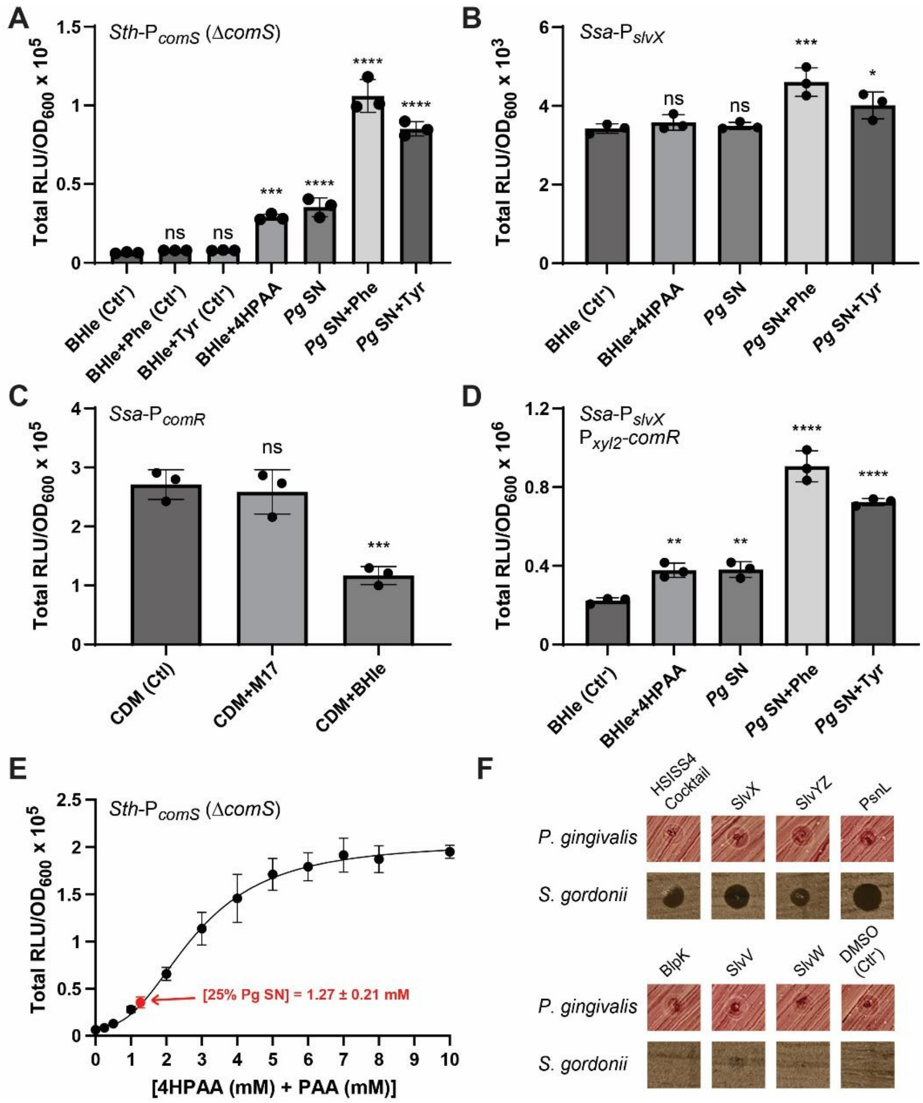

**Fig. S6. Luminescence assays with culture supernatants of *P. gingivalis* and growth inhibition by salvaricins.** (**A**) Luminescence assays (total RLU/$OD_{600}$) of *S. thermophilus* $\mathrm{P}_{comS}$-*luxAB* Δ*comS* with 25% (v/v) of uncultured BHIe medium without and with Phe (15 mM), Tyr (15 mM), or 4HPAA (1 mM) (negative and positive controls), filtered *P. gingivalis* supernatant (*Pg* SN), and *Pg* SN with Phe (15 mM) or Tyr (15 mM) in CDM. (**B**) Luminescence assays of *S. salivarius* $\mathrm{P}_{slvX}$-*luxAB* as reported in panel A. (**C**) Luminescence assays of *S. salivarius* $\mathrm{P}_{comR}$-*luxAB* without (CDM, Ctl) and with 25% (v/v) M17 or BHIe. (**D**) Luminescence assays of *S. salivarius* $\mathrm{P}_{slvX}$-*luxAB* $\mathrm{P}_{xyl2}$-*comR* as reported in panel B. The *comR* expression was induced by adding 0.2% xylose. (**E**) Luminescence assays of *S. thermophilus* $\mathrm{P}_{comS}$-*luxAB* Δ*comS* with increasing concentration (mM) of a mixture of 4HPAA and PAA (ratio 1:1). Each compound of the mixture was individually resuspended in BHIe medium at the indicated concentration. 25% (v/v) of BHIe solutions were incorporated in CDM. The red arrow corresponds to the concentration detected in *Pg* SN. (**F**) Bacteriocin assays with synthetic versions of salivaricins from *S. salivarius* HSISS4 either used either as a cocktail of 6 peptides or tested separately. The broad-spectrum salivaricin PsnL (not produced by HSISS4) was also tested. *P. gingivalis* W83 and *S. gordonii* LMG 17843 were used as indicator strains. DMSO (100%) was used as negative control. In panels A to D, dots, bars, and error bars show biological triplicates, mean values, and standard deviations, respectively. All tested conditions were statistically compared to the control (Ctl) using a one-way ANOVA with Dunnett's test (*, $P < 0.05$; **, $P < 0.01$; ***, $P < 0.001$; ****, $P < 0.0001$; ns, non-significant).

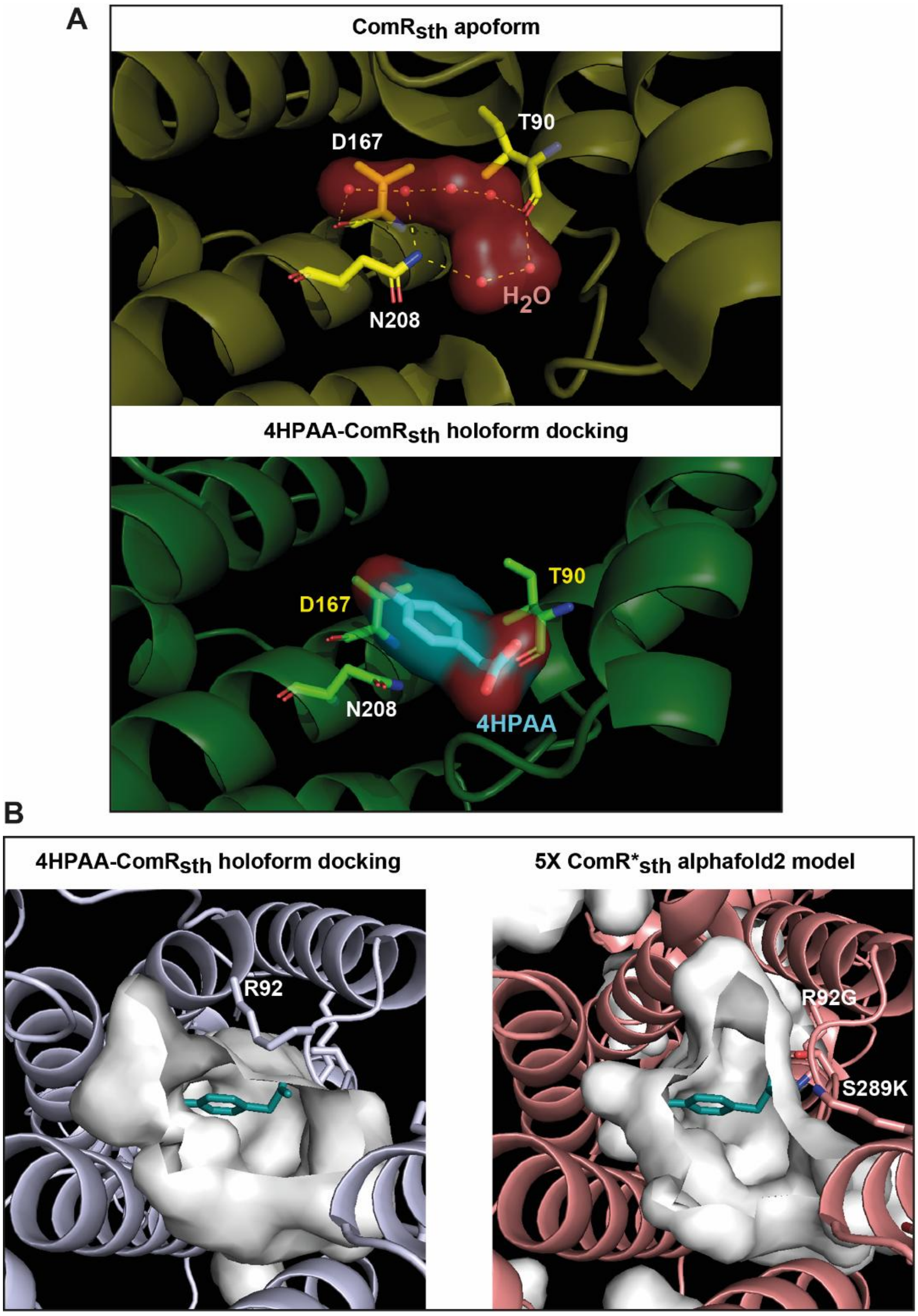

**Fig. S7. Mechanism of ComR$_{Sth}$ activation by 4HPAA.** (**A**) Position of water molecules (red spheres) in the bottom of the XIP-binding pocket of ComR$_{Sth}$ apoform (PDB 5JUF) (top) compared to the predicted docking of 4HPAA (blue) in ComR$_{Sth}$ holoform (PDB 5JUB) (bottom). Four out of six water molecules are interacting with residues T90, D167, and N208 in the apoform (in yellow), which are also predicted to interact with 4HPAA in the holoform (in green). (**B**) Comparison of the size of the 4HPAA-binding pocket between ComR$_{Sth}$ holoform (PDB 5JUB) and 5X ComR*$_{Sth}$ (AlphaFold2 model), mimicking the XIP-binding pocket of ComR$_{Sve}$. The mutation R92G, which recapitulates the absence of 4HPAA induction of 5X ComR*$_{Sth}$, is highlighted. Docked 4HPAA is colored in blue-green.

### SUPPLEMENTAL TABLES

**Table S1. List of tested carbon/nitrogen sources and light emission of reporter strains from Biolog plates PM1 and PM2A.**

| Plate position | Carbon/nitrogen source | Max. light emission (RLU/$OD_{600}$) of reporter strains | | | | | |
|---|---|---|---|---|---|---|---|
| | | $P_{sptA}$ | $P_{slvX}$ | $P_{comX}$ | $P_{comR}$ | $P_{scuR}$ | $P_{sarF}$ |
| | | **Biolog plate PM1** | | | | | |
| A1 | - (control) | 2123 | 2404 | 2036 | 61204 | 46116 | 50886 |
| A2 | L-Arabinose | 2181 | 2966 | 2417 | 74600 | 49243 | 63907 |
| A3 | N-Acetyl-D-Glucosamine | 2808 | 2678 | 1398 | 80847 | 48225 | 81227 |
| A4 | D-Saccharic Acid | 2823 | 2757 | 2177 | 61472 | 39199 | 82843 |
| A5 | Succinic Acid | 2772 | 2689 | 2473 | 33853 | 26325 | 85995 |
| A6 | D-Galactose | 3138 | 3166 | 2540 | 53839 | 43389 | 94963 |
| A7 | L-Aspartic Acid | 3257 | 3315 | 2571 | 61133 | 39693 | 89411 |
| A8 | L-Proline | 3085 | 3270 | 2435 | 53473 | 37290 | 87505 |
| A9 | D-Alanine | 2823 | 2762 | 2646 | 52000 | 30847 | 81173 |
| A10 | D-Trehalose | 3685 | 4586 | 2719 | 130297 | 65608 | 124936 |
| A11 | D-Mannose | 2722 | 2743 | 2560 | 79253 | 42000 | 74934 |
| A12 | Dulcitol | 2929 | 3610 | 2859 | 77354 | 52595 | 56711 |
| B1 | D-Serine | 2365 | 3176 | 2602 | 45272 | 39934 | 48452 |
| B2 | D-Sorbitol | 2638 | 3526 | 2484 | 80326 | 52442 | 59502 |
| B3 | Glycerol | 2689 | 3365 | 2238 | 74669 | 44951 | 87053 |
| B4 | L-Fucose | 2863 | 3155 | 2296 | 42277 | 29051 | 96516 |
| B5 | D-Glucuronic Acid | 3199 | 3341 | 2351 | 44807 | 28077 | 104643 |
| B6 | D-Gluconic Acid | 3369 | 3858 | 2168 | 66472 | 41047 | 106234 |
| B7 | D,L-α-Glycerol-Phosphate | 3306 | 3688 | 2723 | 72425 | 39083 | 107944 |
| B8 | D-Xylose | 3089 | 3377 | 2694 | 40649 | 23812 | 90680 |
| B9 | L-Lactic Acid | 3176 | 3387 | 2750 | 54384 | 29481 | 96626 |
| B10 | Formic Acid | 3764 | 3584 | 2898 | 97610 | 58567 | 117638 |
| B11 | D-Mannitol | 3573 | 4669 | 3222 | 101494 | 64654 | 99699 |
| B12 | L-Glutamic Acid | 2941 | 3897 | 3143 | 75494 | 45905 | 57258 |
| C1 | D-Glucose-6-Phosphate | 2179 | 2986 | 2583 | 56093 | 47341 | 50051 |
| C2 | D-Galactonic Acid-γ-Lactone | 2702 | 3228 | 2291 | 67547 | 31722 | 50165 |
| C3 | D,L-Malic Acid | 2448 | 3175 | 2260 | 72238 | 43739 | 79370 |
| C4 | D-Ribose | 2518 | 3399 | 2363 | 44595 | 28127 | 88980 |
| C5 | Tween 20 | 2279 | 3234 | 2037 | 31798 | 33268 | 37188 |
| C6 | L-Rhamnose | 3257 | 3750 | 2191 | 73177 | 34124 | 101052 |
| C7 | D-Fructose | 3958 | 4882 | 2709 | 107770 | 56871 | 160170 |
| C8 | Acetic Acid | 3395 | 3479 | 2438 | 55052 | 37274 | 108873 |
| C9 | α-D-Glucose | 3163 | 3734 | 2591 | 66594 | 29962 | 97817 |
| C10 | Maltose | 3432 | 4117 | 2742 | 78150 | 52066 | 99570 |
| C11 | D-Melibiose | 3202 | 3972 | 2819 | 78281 | 44659 | 98337 |
| C12 | Thymidine | 3603 | 4762 | 2992 | 101552 | 57317 | 85514 |
| D1 | L-Asparagine | 2347 | 3450 | 2296 | 67344 | 50213 | 52524 |

| | | | | | | | |
|---|---|---|---|---|---|---|---|
| D2 | D-Aspartic Acid | 2078 | 3317 | 2071 | 51272 | 25755 | 25795 |
| D3 | D-Glucosaminic Acid | 2467 | 3174 | 2315 | 59521 | 31876 | 55193 |
| D4 | 1,2-Propanediol | 2667 | 3066 | 2175 | 66516 | 42155 | 84586 |
| D5 | Tween 40 | 4872 | 4731 | 1901 | 109569 | 77337 | 115074 |
| D6 | α-Keto-Glutaric Acid | 3335 | 3864 | 2136 | 86430 | 36720 | 84389 |
| D7 | α-Keto-Butyric Acid | 2987 | 3631 | 2269 | 101492 | 50344 | 107879 |
| D8 | α-Methyl-DGalactoside | 3301 | 4413 | 2125 | 73786 | 44331 | 102009 |
| D9 | α-D-Lactose | 3199 | 3856 | 2473 | 63210 | 26576 | 77197 |
| D10 | Lactulose | 3188 | 3566 | 2628 | 77066 | 43817 | 71485 |
| D11 | Sucrose | 3519 | 7089 | 2811 | 131667 | 62308 | 112802 |
| D12 | Uridine | 3963 | 4038 | 2817 | 80389 | 55898 | 80808 |
| E1 | L-Glutamine | 2679 | 3301 | 2451 | 59682 | 40869 | 57552 |
| E2 | m-Tartaric Acid | 2174 | 3411 | 2234 | 68116 | 30059 | 33562 |
| E3 | D-Glucose-1- Phosphate | 2136 | 3582 | 2193 | 66705 | 31642 | 54482 |
| E4 | D-Fructose-6- Phosphate | 2383 | 3458 | 2224 | 77553 | 41862 | 73811 |
| E5 | Tween 80 | 2799 | 4077 | 2006 | 71396 | 50417 | 75380 |
| E6 | α-Hydroxy Glutaric Acid-γLactone | 3332 | 4126 | 2291 | 99230 | 44930 | 95969 |
| E7 | α-Hydroxy Butyric Acid | 3200 | 3809 | 2314 | 118109 | 59374 | 104361 |
| E8 | ß-Methyl-DGlucoside | 2925 | 4162 | 2072 | 94277 | 51753 | 83392 |
| E9 | Adonitol | 3005 | 4019 | 2563 | 98688 | 27349 | 69112 |
| E10 | Maltotriose | 3218 | 4772 | 2742 | 81891 | 55200 | 93896 |
| E11 | 2-Deoxy Adenosine | 3455 | 4729 | 2683 | 74218 | 44220 | 92733 |
| E12 | Adenosine | 4000 | 4906 | 2969 | 109441 | 57632 | 107015 |
| F1 | Glycyl-L-Aspartic Acid | 2846 | 3449 | 2672 | 45856 | 34555 | 55228 |
| F2 | Citric Acid | 2147 | 3647 | 2398 | 70801 | 29268 | 63500 |
| F3 | myo-Inositol | 2548 | 3772 | 2364 | 79802 | 38961 | 62890 |
| F4 | D-Threonine | 2220 | 3649 | 2504 | 80727 | 52594 | 61438 |
| F5 | Fumaric Acid | 2326 | 4161 | 2155 | 119433 | 61267 | 63419 |
| F6 | Bromo Succinic Acid | 3152 | 5154 | 2669 | 135298 | 67851 | 71555 |
| F7 | Propionic Acid | 2943 | 4438 | 2417 | 140959 | 67944 | 91465 |
| F8 | Mucic Acid | 2725 | 4196 | 2182 | 120628 | 64291 | 66688 |
| F9 | Glycolic Acid | 2473 | 3755 | 2590 | 105097 | 31497 | 69087 |
| F10 | Glyoxylic Acid | 2888 | 4637 | 2695 | 82432 | 48429 | 79381 |
| F11 | D-Cellobiose | 2835 | 4674 | 2651 | 113301 | 57929 | 86011 |
| F12 | Inosine | 3508 | 3699 | 2819 | 82582 | 49635 | 81559 |
| G1 | Glycyl-LGlutamic Acid | 2138 | 3374 | 2454 | 33619 | 27434 | 57850 |
| G2 | Tricarballylic Acid | 4567 | 3738 | 2567 | 50394 | 22224 | 66462 |
| G3 | L-Serine | 2136 | 3569 | 2313 | 80674 | 39019 | 51487 |
| G4 | L-Threonine | 2125 | 3841 | 2527 | 84250 | 52731 | 59314 |
| G5 | L-Alanine | 2190 | 3753 | 2690 | 96781 | 60199 | 43787 |
| G6 | L-Alanyl-Glycine | 2585 | 4007 | 2444 | 106365 | 65784 | 70387 |
| G7 | Acetoacetic Acid | 2609 | 3963 | 2591 | 107873 | 61921 | 69317 |
| G8 | N-Acetyl-ß-DMannosamine | 2284 | 4085 | 2832 | 101557 | 63147 | 58284 |
| G9 | Mono Methyl Succinate | 2644 | 3628 | 2711 | 104966 | 54812 | 45265 |
| G10 | Methyl Pyruvate | 2425 | 3559 | 2646 | 92611 | 49631 | 39440 |

| | | | | | | | |
|---|---|---|---|---|---|---|---|
| G11 | D-Malic Acid | 3141 | 3962 | 2817 | 95966 | 49937 | 83188 |
| G12 | L-Malic Acid | 3303 | 3845 | 3064 | 73873 | 45660 | 82648 |
| H1 | Glycyl-L-Proline | 2949 | 3776 | 2675 | 49521 | 39286 | 83462 |
| H2 | **p-Hydroxy Phenyl Acetic Acid** | 2089 | **130682** | 2539 | 71006 | 18674 | 64206 |
| H3 | **m-Hydroxy Phenyl Acetic Acid** | 2669 | **67956** | 2669 | 94770 | 44674 | 73813 |
| H4 | Tyramine | 2198 | 3421 | 2422 | 74324 | 39416 | 67411 |
| H5 | D-Psicose | 2689 | 4753 | 2619 | 114064 | 73082 | 94717 |
| H6 | L-Lyxose | 2468 | 3494 | 2794 | 64551 | 40640 | 49863 |
| H7 | Glucuronamide | 2643 | 3972 | 2732 | 101659 | 62977 | 67622 |
| H8 | Pyruvic Acid | 2206 | 3810 | 2648 | 95883 | 58876 | 36957 |
| H9 | L-Galactonic Acid-γ-Lactone | 2798 | 3870 | 2738 | 88209 | 27935 | 64854 |
| H10 | D-Galacturonic Acid | 3215 | 3730 | 3039 | 69098 | 44678 | 82353 |
| H11 | Phenylethylamine | 2990 | 3289 | 2798 | 48944 | 29018 | 66543 |
| H12 | 2-Aminoethanol | 3243 | 3476 | 2932 | 30156 | 38638 | 74639 |
| | **Biolog plate PM2A** | | | | | | |
| A1 | - (control) | 2644 | 1438 | 1376 | 30375 | 29502 | 35613 |
| A2 | Chondroitin-Chondroitin Sulfate C | 2769 | 1628 | 1400 | 34043 | 32943 | 53988 |
| A3 | α-Cyclodextrin | 2784 | 1577 | 1444 | 89136 | 63369 | 66016 |
| A4 | ß-Cyclodextrin | 2865 | 1626 | 1483 | 68433 | 64073 | 49773 |
| A5 | γ-Cyclodextrin | 3114 | 1632 | 1531 | 47445 | 46408 | 34084 |
| A6 | Dextrin | 3069 | 1667 | 1623 | 38254 | 41289 | 31418 |
| A7 | Gelatin | 2985 | 1772 | 1535 | 48174 | 38509 | 34774 |
| A8 | Glycogen | 3000 | 1852 | 1688 | 52654 | 42491 | 34160 |
| A9 | Inulin | 2594 | 1447 | 1424 | 37954 | 40221 | 34764 |
| A10 | Laminarin | 3061 | 1862 | 1793 | 37610 | 34286 | 31330 |
| A11 | Mannan | 2864 | 2084 | 469 | 33178 | 34821 | 31562 |
| A12 | Pectin | 2830 | 2031 | 1846 | 51244 | 49311 | 28115 |
| B1 | N-Acetyl-D-Galactosamine | 2783 | 2016 | 1833 | 27758 | 25659 | 29314 |
| B2 | N-Acetyl-Neuraminic Acid | 2857 | 1679 | 1617 | 37620 | 31197 | 43745 |
| B3 | ß-D-Allose | 2934 | 2048 | 1691 | 54823 | 42932 | 37881 |
| B4 | Amygdalin | 2977 | 1782 | 1603 | 41323 | 32522 | 24915 |
| B5 | D-Arabinose | 3030 | 1846 | 1650 | 55380 | 45648 | 27182 |
| B6 | D-Arabitol | 3131 | 1992 | 1777 | 64094 | 53166 | 39095 |
| B7 | L-Arabitol | 3069 | 2000 | 1846 | 61802 | 49851 | 38546 |
| B8 | Arbutin | 2896 | 1917 | 1702 | 58619 | 43798 | 28626 |
| B9 | 2-Deoxy-DRibose | 3478 | 2598 | 2564 | 35599 | 35323 | 28370 |
| B10 | i-Erythritol | 3180 | 2135 | 1936 | 53297 | 44177 | 33000 |
| B11 | D-Fucose | 3667 | 2157 | 1941 | 53446 | 41887 | 33991 |
| B12 | 3-0-ß-DGalactopyranosylD-Arabinose | 3209 | 2626 | 2507 | 42586 | 24845 | 36132 |
| C1 | Gentiobiose | 2713 | 1805 | 1550 | 27052 | 25874 | 22148 |
| C2 | L-Glucose | 2649 | 1567 | 1437 | 34061 | 31589 | 26552 |

| | | | | | | | |
|---|---|---|---|---|---|---|---|
| C3 | Lactitol | 2767 | 1662 | 1191 | 42584 | 34952 | 38070 |
| C4 | D-Melezitose | 2856 | 1654 | 1377 | 57688 | 40411 | 38762 |
| C5 | Maltitol | 2924 | 1891 | 1528 | 61259 | 56510 | 36472 |
| C6 | α-Methyl-DGlucoside | 2938 | 2122 | 1472 | 70652 | 62717 | 48624 |
| C7 | ß-Methyl-DGalactoside | 2992 | 1984 | 1450 | 65615 | 52957 | 49994 |
| C8 | 3-Methyl Glucose | 2962 | 1912 | 1674 | 71782 | 53164 | 39715 |
| C9 | ß-Methyl-DGlucuronic Acid | 2992 | 2078 | 1822 | 34502 | 44956 | 32418 |
| C10 | α-Methyl-DMannoside | 3305 | 2000 | 1949 | 58977 | 42599 | 31704 |
| C11 | ß-Methyl-DXyloside | 2992 | 2048 | 2119 | 52342 | 44601 | 34261 |
| C12 | Palatinose | 3626 | 2440 | 2364 | 41236 | 34715 | 37719 |
| D1 | D-Raffinose | 2529 | 1810 | 1730 | 35029 | 30479 | 7686 |
| D2 | Salicin | 2939 | 1284 | 1367 | 9249 | 17039 | 17284 |
| D3 | Sedoheptulosan | 2649 | 1349 | 1315 | 26836 | 36244 | 28735 |
| D4 | L-Sorbose | 2809 | 1500 | 1419 | 52929 | 50099 | 47056 |
| D5 | Stachyose | 2689 | 1489 | 1441 | 63702 | 55393 | 41416 |
| D6 | D-Tagatose | 2550 | 1865 | 1452 | 65009 | 54230 | 50399 |
| D7 | Turanose | 2481 | 1887 | 1588 | 64158 | 58525 | 56929 |
| D8 | Xylitol | 2802 | 1939 | 1835 | 40442 | 41731 | 29478 |
| D9 | N-Acetyl-DGlucosaminitol | 3084 | 2087 | 1815 | 40727 | 40942 | 38016 |
| D10 | γ-Amino Butyric Acid | 2909 | 2024 | 1880 | 44477 | 33705 | 33274 |
| D11 | δ-Amino Valeric Acid | 3231 | 2215 | 1913 | 34853 | 29682 | 29361 |
| D12 | Butyric Acid | 3153 | 2273 | 2319 | 46464 | 40392 | 45795 |
| E1 | Capric Acid | 2933 | 1887 | 1899 | 4604 | 3549 | 3612 |
| E2 | Caproic Acid | 2779 | 2147 | 1530 | 16000 | 18545 | 26800 |
| E3 | Citraconic Acid | 2626 | 1603 | 1418 | 33151 | 34424 | 24533 |
| E4 | Citramalic Acid | 2740 | 1638 | 1489 | 42361 | 34863 | 51919 |
| E5 | D-Glucosamine | 2629 | 1686 | 1504 | 27234 | 36889 | 44021 |
| E6 | 2-Hydroxy Benzoic Acid | 2924 | 1895 | 1590 | 129479 | 51654 | 66672 |
| E7 | 4-Hydroxy Benzoic Acid | 3000 | 1746 | 1551 | 115839 | 54964 | 56791 |
| E8 | ß-Hydroxy Butyric Acid | 3382 | 1014 | 1647 | 48207 | 42923 | 54616 |
| E9 | Glycolic Acid | 2948 | 1968 | 1779 | 22596 | 40594 | 45340 |
| E10 | α-Keto-Valeric Acid | 2253 | 2143 | 1947 | 40037 | 33873 | 33690 |
| E11 | Itaconic Acid | 3208 | 2148 | 1882 | 38472 | 36641 | 28165 |
| E12 | 5-Keto-DGluconic Acid | 3235 | 2421 | 2362 | 47699 | 39307 | 9109 |
| F1 | D-Lactic Acid Methyl Ester | 2730 | 1976 | 1621 | 27201 | 27041 | 10602 |
| F2 | Malonic Acid | 2646 | 1563 | 1570 | 30664 | 27333 | 37443 |
| F3 | Melibionic Acid | 3735 | 1532 | 1513 | 33349 | 31846 | 39308 |
| F4 | Oxalic Acid | 2463 | 1477 | 1439 | 29872 | 28360 | 52704 |
| F5 | Oxalomalic Acid | 2607 | 1518 | 1649 | 39304 | 37194 | 56888 |
| F6 | Quinic Acid | 2748 | 1468 | 1541 | 50146 | 43395 | 70979 |
| F7 | D-Ribono-1,4- Lactone | 2823 | 1706 | 1569 | 54406 | 44485 | 66223 |
| F8 | Sebacic Acid | 2977 | 1890 | 1777 | 91792 | 44614 | 68545 |
| F9 | Sorbic Acid | 2809 | 2083 | 1914 | 12386 | 14783 | 25060 |
| F10 | Succinamic Acid | 2856 | 1946 | 1841 | 39561 | 33783 | 47504 |
| F11 | D-Tartaric Acid | 3046 | 1985 | 2051 | 36905 | 38576 | 34115 |

| | | | | | | | |
|---|---|---|---|---|---|---|---|
| F12 | L-Tartaric Acid | 3618 | 2282 | 2378 | 50044 | 40313 | 43925 |
| G1 | Acetamide | 2842 | 1754 | 1697 | 25292 | 19576 | 22821 |
| G2 | L-Alaninamide | 2733 | 1677 | 1696 | 23906 | 28247 | 40446 |
| G3 | N-Acetyl-LGlutamic Acid | 2591 | 1597 | 1763 | 13575 | 24902 | 36821 |
| G4 | L-Arginine | 2629 | 1643 | 1630 | 24678 | 30397 | 54346 |
| G5 | Glycine | 2604 | 1711 | 1606 | 20065 | 28798 | 61027 |
| G6 | L-Histidine | 2977 | 1819 | 1688 | 36381 | 34614 | 64974 |
| G7 | L-Homoserine | 2659 | 2024 | 1704 | 41798 | 38957 | 68118 |
| G8 | Hydroxy-LProline | 2970 | 1922 | 1956 | 35982 | 31373 | 52413 |
| G9 | L-Isoleucine | 2923 | 2192 | 1779 | 16205 | 27645 | 54828 |
| G10 | L-Leucine | 3113 | 2221 | 1905 | 29591 | 23387 | 48509 |
| G11 | L-Lysine | 2992 | 2102 | 2022 | 46728 | 38761 | 36467 |
| G12 | L-Methionine | 3344 | 2560 | 2356 | 59804 | 48263 | 42720 |
| H1 | L-Ornithine | 2718 | 1739 | 1694 | 34980 | 33415 | 19611 |
| H2 | L-Phenylalanine | 2939 | 1721 | 1837 | 22556 | 24194 | 21018 |
| H3 | L-Pyroglutamic Acid | 2941 | 1897 | 1734 | 26722 | 33415 | 48760 |
| H4 | L-Valine | 2881 | 1953 | 1750 | 31249 | 35951 | 50057 |
| H5 | D,L-Carnitine | 2835 | 1806 | 1854 | 30934 | 33058 | 56784 |
| H6 | Sec-Butylamine | 2858 | 2180 | 1894 | 26886 | 35182 | 55409 |
| H7 | D,L-Octopamine | 3233 | 2114 | 1924 | 33881 | 28608 | 47641 |
| H8 | Putrescine | 3105 | 2138 | 1943 | 23851 | 19150 | 58247 |
| H9 | Dihydroxy Acetone | 4490 | 2809 | 2699 | 20653 | 15027 | 27378 |
| H10 | 2,3-Butanediol | 3173 | 2231 | 2096 | 40179 | 24338 | 36664 |
| H11 | 2,3-Butanedione | 3160 | 2365 | 2207 | 49597 | 30890 | 33867 |
| H12 | 3-Hydroxy-2- Butanone | 3542 | 2346 | 2300 | 44947 | 35926 | 27807 |

**Table S2. List of organic acids and amino acids tested in this study.**

| Common name | Systematic IUPAC | Abbreviation | Supplier |
|---|---|---|---|
| 4-hydroxy phenylacetic acid | (4-Hydroxyphenyl)acetic acid | 4HPAA | Thermo Fisher Scientific |
| 3-Hydroxy phenylacetic acid | 2-(3-Hydroxyphenyl)acetic acid | 3HPAA | Merck |
| Tyramine | 4-(2-Aminoethyl)phenol | | Thermo Fisher Scientific |
| Methyl 4-hydroxyphenylacetate | Methyl 2-(4-hydroxyphenyl)acetate | Methyl-4HPAA | Thermo Fisher Scientific |
| D-(+)-3-Phenyllactic acid | (R)-2-Hydroxy-3-phenylpropionic acid | | Thermo Fisher Scientific |
| (R)-(-)-Mandelic Acid | (2R)-2-Hydroxy-2-phenylacetic acid | | Thermo Fisher Scientific |
| Phenylacetic acid | 2-Phenylethanoic acid | PAA | Merck |
| 3-Fluoro-4-hydroxyphenylacetic acid | 2-(3-Fluoro-4-hydroxyphenyl)acetic acid | 3-Fluoro-4HPAA | Thermo Fisher Scientific |
| Pentafluoro phenylacetic acid | 2,3,4,5,6-Pentafluorophenylacetic acid | Pentafluoro-PAA | Thermo Fisher Scientific |
| Homovanillic acid | (4-Hydroxy-3-methoxyphenyl)acetic acid | | Thermo Fisher Scientific |
| 4-Hydroxy-3,5-dimethoxyphenylacetic acid | 2-(4-Hydroxy-3,5-dimethoxyphenyl)acetic acid | 4-Hydroxy-3,5-dimethoxy-PAA | Thermo Fisher Scientific |
| 1-Naphthylacetic acid | 2-(Naphthalen-1-yl)acetic acid | | Thermo Fisher Scientific |
| Indole-3-acetic acid | 2-(1H-indol-3-yl)acetic acid | IAA | Thermo Fisher Scientific |
| 3-Methylvaleric acid | 3-Methylpentanoic acid | 3-MVA | Thermo Fisher Scientific |
| 4-Methylvaleric acid | 4-Methylpentanoic acid | 4-MVA | Thermo Fisher Scientific |
| Isovaleric acid | 3-Methylbutanoic acid | | Thermo Fisher Scientific |
| 4-Imidazolacetic acid | Butyl 2-(1*H*-imidazol-5-yl)acetate | | Thermo Fisher Scientific |
| Propionic acid | Propionic acid | | Thermo Fisher Scientific |
| Tyrosine | L-2-Amino-3-(4-hydroxyphenyl)propanoic acid | | Thermo Fisher Scientific |
| Phenylalanine | (*S*)-2-Amino-3-phenylpropanoic acid | | Thermo Fisher Scientific |
| Tryptophan | (2*S*)-2-Amino-3-(1*H*-indol-3-yl)propanoic aci | | Thermo Fisher Scientific |
| Histidine | 2-Amino-3-(1*H*-imidazol-4-yl)propanoic acid | | Thermo Fisher Scientific |
| Alanine | 2-Aminopropanoic acid | | Thermo Fisher Scientific |
| Valine | 2-Amino-3-methylbutanoic acid | | Thermo Fisher Scientific |
| Isoleucine | (2S,3S)-2-Amino-3-methylpentanoic acid | | Thermo Fisher Scientific |
| Leucine | 2-Amino-4-methylpentanoic acid | | Thermo Fisher Scientific |

### Table S3. List of strains and plasmids.

| Strain/plasmid | Genotype/description | Resistance[a] | Source |
|---|---|---|---|
| **Strains** | | | |
| *Escherichia coli* | | | |
| *E. coli* TOP10 | $F^-$ *mcrA* Δ(*mrr-hsdRMS mcrBC*) φ80*lacZ*Δ*M15* Δ*lacX74 recA1 araD139* Δ(*ara-leu*) 7697 *galU galK rpsL* ($Str^r$) *endA1 nupG λ-* | - | Invitrogen, CA |
| *Streptococcus thermophilus* | | | |
| LMD-9 | Wild-type milk isolate | - | ATCC[b] |
| LF121 | LMD-9 (*blpD-blpX*)::$P_{comS}$-*luxAB* | - | (1) |
| LF134 | LF121 *ΔcomS*::$P_{32}$-*cat* | $Cm^R$ | (1) |
| LF135 | LF121 *ΔcomR::*$P_{32}$-*cat* | $Cm^R$ | (1) |
| LF138 | LF121 *ΔamiA1-amiF::*$P_{32}$-*cat* | $Cm^R$ | L. Fontaine, Lab. collection |
| LF149 | LF121 *comR::comR*$_{T90A}$, *ΔcomS::*$P_{32}$-*cat* | $Cm^R$ | (2) |
| LF152 | LF121 *comR::comR*$_{K100A}$, *ΔcomS::*$P_{32}$-*cat* | $Cm^R$ | (2) |
| LF153 | LF121 *comR::comR*$_{F171A\text{-}Y174A}$, *ΔcomS*::$P_{32}$-*cat* | $Cm^R$ | (2) |
| LL1 | LF121 *comR::comR*$_{Sve}$, *ΔcomS::*$P_{32}$-*cat* | $Cm^R$ | (3) |
| LL10 | LF121 *comR::comR*$_{R92G\text{-}P94K}$, *ΔcomS::*$P_{32}$-*cat; L6** | $Cm^R$ | (3) |
| LL11 | LF121 *comR::comR*$_{V201A\text{-}V205A}$, *ΔcomS::*$P_{32}$-*cat; α12** | $Cm^R$ | (3) |
| LL12 | LF121 *comR::comR*$_{S248G}$, *ΔcomS::*$P_{32}$-*cat; α14** | $Cm^R$ | (3) |
| LL13 | LF121 *comR::comR*$_{S289K\text{-}I290T}$, *ΔcomS::*$P_{32}$-*cat; CAP** | $Cm^R$ | (3) |
| LL15 | LF121 *comR::comR*$_{V205A}$, *ΔcomS::*$P_{32}$-*cat* | $Cm^R$ | (3) |
| LL19 | LF121 *comR::comR*$_{R92G\text{-}V205A\text{-}S248G\text{-}S289K\text{-}I290T}$, *ΔcomS::*$P_{32}$-*cat* | $Cm^R$ | (3) |
| LL40 | LF121 *comR::comR*$_{F171A}$, *ΔcomS::*$P_{32}$-*cat* | $Cm^R$ | (4) |
| LL42 | LF121 *comR::comR*$_{Y174A}$, Δ*comS*::$P_{32}$-*cat* | $Cm^R$ | (4) |
| LL100 | LF121 *comR::comR*$_{R92G}$, *ΔcomS::*$P_{32}$-*cat* | $Cm^R$ | (3) |
| LL101 | LF121 *comR::comR*$_{P94K}$, *ΔcomS::*$P_{32}$-*cat* | $Cm^R$ | (3) |
| LL102 | LF121 *comR::comR*$_{S289K}$, *ΔcomS::*$P_{32}$-*cat* | $Cm^R$ | (3) |
| LL103 | LF121 *comR::comR*$_{I290T}$, *ΔcomS::*$P_{32}$-*cat* | $Cm^R$ | (3) |
| *Streptococcus salivarius* | | | |
| HSISS4 | Wild-type gastro-intestinal tract isolate | - | (5) |
| JM1001 | HSISS4 Δ*comR* | - | (6) |
| JM1013 | HSISS4 Δ*slv5* | - | (6) |
| JM1016 | HSISS4 *tRNA*$^{ser}$*::*$P_{xyl2}$-*comR-spc* | $Spec^R$ | (6) |
| JM1020 | HSISS4 *tRNA*$^{Thr}$*::*$P_{comX}$-*luxAB-cat* | $Cm^R$ | (6) |
| JM1027 | HSISS4 *tRNA*$^{Thr}$*::*$P_{slvX}$-*luxAB-cat* | $Cm^R$ | (6) |
| JM1100 | HSISS4 *tRNA*$^{Thr}$*::*$P_{sptA}$-*luxAB-cat* | $Cm^R$ | (7) |
| JM1101 | HSISS4 *tRNA*$^{Ser}$*::*$P_{32}$-*scuR-spc* (*scuR*$^{++}$) | $Spec^R$ | (7) |

| | | | |
|---|---|---|---|
| JM1118 | HSISS4 Δ*scuR-sarF::erm* | $Ery^R$ | (7) |
| JM1192 | JM1027 Δ*scuR-sarF::erm* | $Cm^R$ & $Ery^R$ | (7) |
| JM1175 | JM1100 Δ*scuR::erm* | $Ery^R$ | (7) |
| JM1300 | JM1027 Δ*comS::erm* | $Cm^R$ & $Ery^R$ | This work |
| JM1301 | JM1027 Δ*comR::erm* | $Cm^R$ & $Ery^R$ | This work |
| JM1302 | HSISS4 $tRNA^{Thr}$::$P_{comR}$-*luxAB-cat* | $Cm^R$ | This work |
| JM1303 | HSISS4 $tRNA^{Thr}$::$P_{scuR}$-*luxAB-cat* | $Cm^R$ | This work |
| JM1304 | HSISS4 $tRNA^{Thr}$::$P_{sarF}$-*luxAB-cat* | $Cm^R$ | This work |
| JM1305 | JM1027 $tRNA^{Ser}$::$P_{xyl2}$-*comR-cat* | $Cm^R$ | This work |
| *Lactococcus lactis* | | | |
| IL1403 | Laboratory strain | - | (8) |
| *Porphyromonas gingivalis* | | | |
| W83 (ATCC BAA-308) | Wild-type clinical isolate | - | ATCC[b] |
| *Streptococcus gordonii* | | | |
| LMG 17843 | Wild-type isolate from human oral cavity | | BCCM[c] |
| **Plasmids** | | | |
| pBAD-*comR*$_{Sth}$-*strep* | pBADHisA derivative containing the translation fusion $P_{ara}$BAD-*comR*$_{Sth}$-*strep* | $Ap^R$ | (1) |
| pBAD-*comR*$_{Sve}$-*strep* | pBADHisA derivative containing the translation fusion $P_{ara}$BAD-*comR*$_{Sve}$-*strep* | $Ap^R$ | (3) |
| pGIUD0855*ery* | pUC18 derivative containing the *erm* gene | $Ery^R$ | (9) |
| pJIM*cat* | pJIM4900 derivative with a *cat* cassette | $Cm^R$ | (7) |

[a]$Ap^R$, $Cm^R$, $Ery^R$, and $Spec^R$; resistance to ampicillin, chloramphenicol, erythromycin, and spectinomycin, respectively.
[b]ATCC, American Type Culture Collection, Rockville, MD.
[c]BCCM, Belgian Co-ordinated Collections of Micro-organisms

**Table S4. List of oligonucleotides used in this study.**

**Construction of *S. salivarius* strains**

| Names | Sequence (5' to 3') |
|---|---|
| Uplox66 | TAAGGAAGATAAATCCCATAAGG |
| DNlox71 | TTCACGTTACTAAAGGGAATGTA |
| lox66-ery | TAAGGAAGATAAATCCCATAAGGTACCTAATAATTTATCTACATTCC |
| lox71-ery | TTCACGTTACTAAAGGGAATGTAAAATGATACACCAATCAGTGC |
| UF_tRNAthr | TGTCAAAGGATTAGGAAAAC |
| UR_tRNAthr | TTGATTTATACCTCTCAATTT |
| DF_tRNAthr | AAATCAACCTCTTTGAACATA |
| DR_tRNAthr | AAAAAAGAATTCATTCATGATGAGCGGGTTCGTGAGA |
| F_luxAB_ATG | ATGAAATTTGGAAACTTTTTGC |
| R_cat_tRNAthr | TATGTTCAAAGAGGTTGATTTCACGTTACTAAAGGGAATGTA |
| UFcomRJIM-SS1-4 | GCAGTACCACTCTATGCTAAATTTGCCAACTTTGA |
| URcomRJIM-SS1-4 | CCTTATGGGATTTATCTTCCTTAGAGACACTCCTTTATTTC |
| DFcomRJIM-SS1-4 | TACATTCCCTTTAGTAACGTGAAAAATGGTGGTGACATAAA |
| DRcomRJIM-SS1-4 | TGACGTGATTTCACCAGTACGACGTGAACTAAAGA |
| Up_comR SS1-4 | TTGCTTACAGTTGCTATGGT |
| Down_comR_SS1-4 | TCATCACAATGGTCACATCT |
| UF_PcomR_luxAB | TAATTGAGGAGGTCTATGAG |
| UR_comS | CCTTATGGGATTTATCTTCCTTATAAAACTCCTTTTAACTGTTAG |
| DF_comS | TACATTCCCTTTAGTAACGTGAATAATAAGGAGTCACCATGTC |
| F_comR | CTAGAGGAGGAATTTAGATGAACATAAAAGACAGCATTG |
| Down_PcomS_JIMSS1-4 | GACAAAGTAGTCAAGACCGT |
| F_PrggD_tRNAthr | AAATTGAGAGGTATAAATCAATGCTATAATTTTCATCATCG |
| F_PrggC_tRNAthr | AAATTGAGAGGTATAAATCAATCCTATTTATAACACTGACC |
| R_PcomR_luxAB_ATG | GCAAAAAGTTTCCAAATTTCAAGAGACACTCCTTTATTT |
| R_PrggD_luxAB_ATG | GCAAAAAGTTTCCAAATTTCATACATAATTCCTTATGATTT |
| R_PrggC_luxAB_ATG | GCAAAAAGTTTCCAAATTTCATCTGTATTTCCCTTGAG |

**Transformation assays**

| Name | Sequence (5' to 3') | Usage | Source |
|---|---|---|---|
| UF_rggC_JIM | AAAACTGCAAGTAGAGTCGC CGAATTAGAA | Linear DNA fragment amplified from strain JM1175 (Δ*scuR::erm*) and used as donor DNA | (7) |
| DR_rggD_SS | TAGCTTCATTCATGTCATGTG TCGTCAAAA | | (7) |

**EMSAs**

| Name | Sequence (5' to 3') | Usage | Source |
|---|---|---|---|
| Cy3-Fw.ComSboxSth.com. direct | ATAGAAATGGTGGTGACATA AATGTCACTATTTTTTTTAG | Probe Cy3-ComR box of $P_{comS}$ from strain LMD-9 | (1) |
| Rv.ComSboxSth.com.direct | CTAAAAAAAATAGTGACATTT ATGTCACCACCATTTCTAT | | (1) |

**Table S5. Overlapping and cloning PCR sub-fragments**

| PCR | Primer 1 | Primer 2 |
|---|---|---|
| *Erm* cassette amplification | lox66-ery | lox71-ery |
| *luxAB-cat* amplification | F_luxAB_ATG | R_cat_tRNAthr |
| Upstream homologous region of $tRNA_{thr}$ locus | UF_tRNAthr | UR_tRNAthr |
| Downstream homologous region of $tRNA_{thr}$ locus | DF_tRNAthr | DR_tRNAthr |
| Upstream homologous region of *comR* gene | UFcomRJIM-SS1-4 | URcomRJIM-SS1-4 |
| Downstream homologous region of *comR* gene | DFcomRJIM-SS1-4 | DRcomRJIM-SS1-4 |
| Diagnostic PCR for *comR* deletion | Up_comR SS1-4 | Down_comR_SS1-4 |
| Upstream homologous region of *comS* gene | UF_PcomR_luxAB | UR_comS |
| Downstream homologous region of *comS* gene | DF_comS | DRcomRJIM-SS1-4 |
| Diagnostic PCR for *comS* deletion | F_comR | Down_comS_SS1-4 |
| Promoter of *comR* for *luxAB* fusion | F_PcomR_tRNAthr | R_PcomR_luxAB_ATG |
| Promoter of *scuR* for *luxAB* fusion | F_PrggD_tRNAthr | R_PrggD_luxAB_ATG |
| Promoter of *sarF* for *luxAB* fusion | F_PrggC_tRNAthr | R_PrggC_luxAB_ATG |
